# Structure of a Stand-Alone Homodimeric NRPS Condensation Domain Reveals Occlusion of the Canonical Carrier-Protein Interface

**DOI:** 10.64898/2026.04.24.720670

**Authors:** Jitendra Singh, Thomas D. Grant, Andrew M. Gulick

## Abstract

Fatty acid amides (FAAs) produced by gut-resident bacteria act as potent modulators of host G protein–coupled receptor signaling, yet the enzymatic mechanisms underlying their biosynthesis remain poorly understood. In many bacteria from the gut microbiome, including *Coprococcus eutactus*, FAA production is mediated by a nonribosomal peptide synthetase–like pathway that includes OaaC, a free-standing condensation domain that catalyzes amide bond formation between acyl carrier protein (ACP) tethered fatty acids and small-molecule amine acceptors. Here, we combine structural, biophysical, biochemical, and evolutionary analyses to interrogate the molecular basis of OaaC function. Solution scattering and X-ray crystallography reveal that OaaC adopts an atypical homodimeric architecture that occludes the canonical ACP-binding surface and donor access pathways. Mass photometry demonstrates that this homodimer is stable in the absence of substrates and is insensitive to free fatty acids, free amines, and apo-ACP. In contrast, holo or acyl-loaded OaaACP selectively destabilizes the homodimer forming the OaaC-OaaACP complex population. LC–MS reconstitution assays confirm that OaaC catalyzes fatty acid amide formation in vitro and can utilize acyl donors spanning multiple chain lengths and saturation states. Phylogenetic and sequence analyses place FAA-associated condensation domains in a distinct clade most closely related to starter condensation domains and reveal a conserved noncanonical active site motif that differentiates them from PCP-dependent NRPS condensation domains. Together, these findings support a model in which OaaC activity is regulated through substrate-dependent modulation of oligomeric state, providing a model framework for understanding FAA biosynthesis in gut microbes and expanding the known functional diversity of NRPS condensation domains.

## Introduction

Nonribosomal peptide synthetases (NRPSs) are modular enzymatic assembly lines that catalyze the biosynthesis of a wide range of peptide and peptide-like natural products, many of which possess important biological and clinical activities (1–5). Canonical NRPS systems are typically encoded within biosynthetic gene clusters (BGCs) and organized as large, multidomain proteins composed of adenylation (A), peptidyl carrier protein (PCP), condensation (C), and, in many cases, thioesterase (TE) domains (6–8). In these systems (Fig. 1A), A domains activate amino acid substrates as aminoacyl adenylates and transfer them to the phosphopantetheine arm of adjacent PCP domains (9). The PCP domains shuttle intermediates between catalytic centers, where C domains catalyze peptide bond formation, and TE domains release the final product (10–12). Organized as multidomain assembly lines, the NRPS enzymes catalyze the formation of the growing peptide that is ultimately released through hydrolysis, aminolysis, cyclization or reduction.

**Figure 1.**
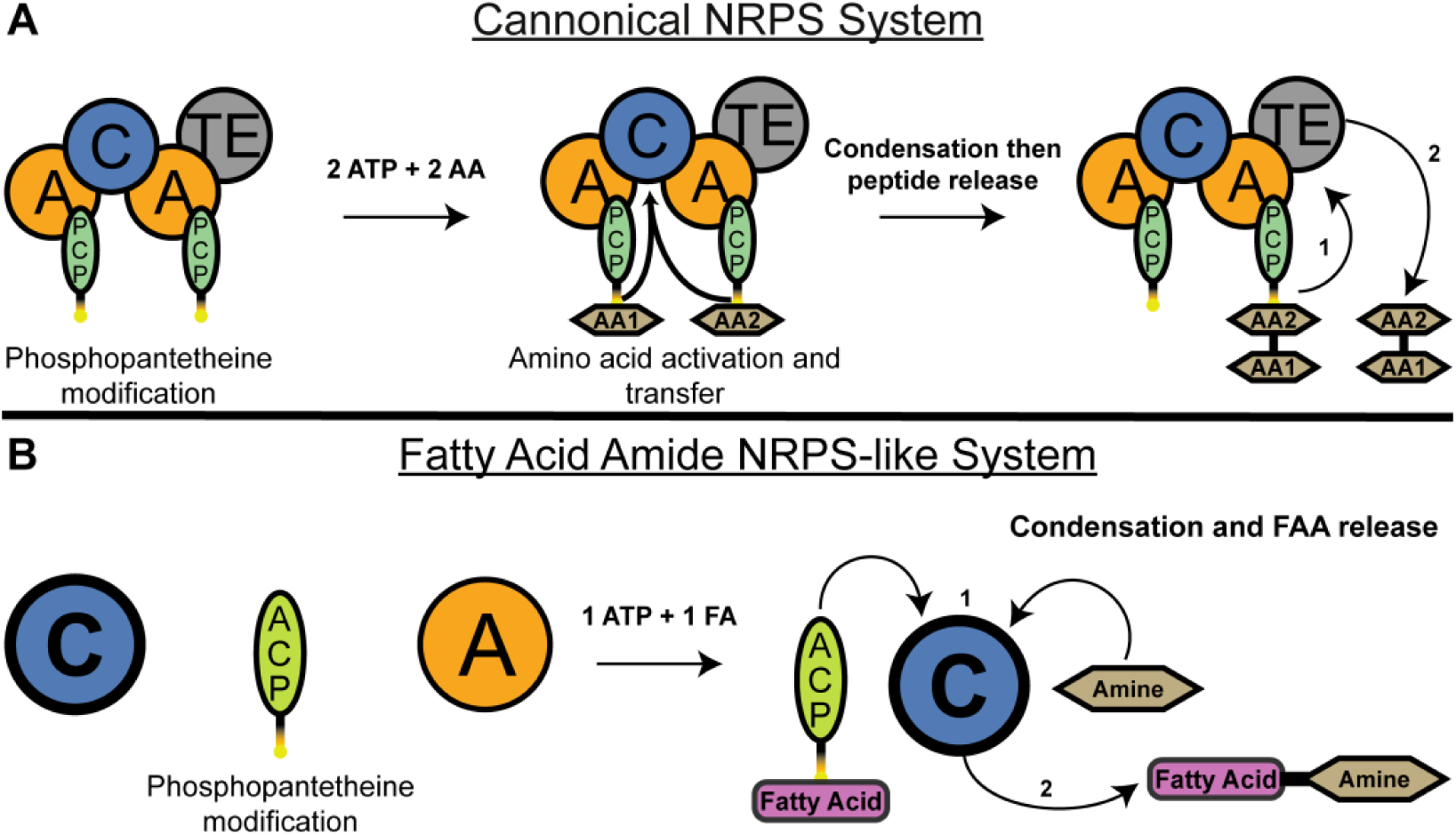
Canonical NRPS and fatty acid amide NRPS-like protein pathways. **A**. A canonical two module NRPS system utilizes a protein with two adenylation (A) domains that activate amino acid substrates, two peptidyl carrier protein (PCP) domains that transport the activated amino acids, a condensation (C) domain that catalyzes peptide bond formation, and a thioesterase (TE) domain for product release. Following phosphopantetheinylation of the carrier protein domains by a phosphopantetheinyl transferase, each A domain separately activates its cognate amino acid substrate using ATP and transfers it to the adjacent PCP domain. The PCP-bound amino acids then meet at the C domain for condensation to form a peptide bond. The dipeptide product is subsequently transferred to the catalytic serine or cysteine within the TE domain (1) and then hydrolyzed (2) from the NRPS complex by the TE domain and released. **B**, the fatty acid amide NRPS-like systems employ a simplified architecture with separate C, A, and acyl carrier protein (ACP) domains. The A domain activates a fatty acid substrate using one ATP molecule and transfers it to the *holo*-ACP domain. The acyl-ACP then delivers the fatty acid to the C domain (1). Within the C domain, the fatty acid undergoes condensation with a small molecule amine substrate, resulting in fatty acid amide product formation and release (2).

Because of their modularity and catalytic versatility, NRPS systems have been the focus of extensive engineering efforts aimed at generating novel natural products with altered or improved biological properties (13–17). Approaches such as domain swapping, substrate specificity reprogramming, and directed evolution have enabled the biosynthesis of modified lipopeptides and small peptides with diverse pharmacological activities (18–20). However, engineered NRPS systems frequently exhibit reduced product yields relative to native assembly lines. These limitations are often attributed to impaired interdomain communication, inefficient substrate transfer, or altered protein–protein interactions between engineered and downstream domains (19, 21).

Among NRPS domains, condensation domains represent a particularly important determinant of pathway efficiency. They catalyze the formation of a peptide bond between two proteogenic amino acids through precise substrate orientation. Canonical C domains consist of N- and C-terminal lobes that adopt a chloramphenicol acetyltransferase-like fold, forming a characteristic V-shaped architecture (22–24). The active site lies at the interface of the two lobes and typically contains a conserved HHxxxDG motif essential for catalysis (25). Structural and biochemical studies have demonstrated that productive condensation depends not only on substrate chemistry but also on precise positioning and orientation of donor and acceptor PCP domains within the active site (26, 27). Recent high-resolution structures of intact NRPS modules captured in pre- and post-condensation states have provided direct evidence that transient PCP–C domain interactions govern catalytic competence (27). However, these insights are derived almost exclusively from canonical, multidomain NRPS assembly lines and do not address how condensation domains function outside this architectural framework.

Recent studies of the human gut microbiome have identified an emerging class of NRPS-like systems that deviate from canonical assembly-line organization. Chang and colleagues discovered multiple biosynthetic gene clusters in gut-resident *Clostridia* that encode enzymes responsible for the biosynthesis of fatty acid amides (FAAs), a class of small molecules that are either identical to, or structural mimics of, endogenous human signaling lipids (28). These FAAs activate several host G protein–coupled receptors (GPCRs) involved in immune signaling, metabolism, and inflammation, linking microbial secondary metabolism to host physiology.

Unlike canonical NRPSs, FAA-producing NRPS systems are composed of discrete, free-standing proteins rather than large multidomain proteins. These systems (Fig. 1B) typically include a stand-alone adenylation domain that activates a fatty acid substrate and transfers it to a separate acyl carrier protein (ACP), and a free-standing condensation domain that catalyzes amide bond formation between the acyl chain and a small-molecule amine acceptor (28). A recent analysis examines the enrichment or depletion of these FAAs in Crohn’s disease (CD) and ulcerative colitis (UC), illustrating that while some FAA encoding biosynthetic gene clusters are enriched in CD and UC, others, including a system for the production of oleoyl aminovaleric acid, are less well represented in the microbiome of disease patients(29).

FAA NRPS-like condensation domains exhibit several features that distinguish them from canonical NRPS C domains. First, they lack an acceptor PCP domain and instead utilize freely diffusing small-molecule amines as acceptor substrates. In this regard, they are similar to terminating condensation domains that transfer the final peptide to small molecule amine acceptor (30, 31). Second, they display high selectivity for specific amines while excluding proteogenic and nonproteogenic amino acids. Third, sequence analyses indicate that these domains frequently harbor a noncanonical DHxxxDS catalytic motif, rather than the HHxxxDG motif characteristic of canonical C domains (28). Together, these observations suggest that FAA condensation domains have evolved distinct structural and mechanistic strategies for substrate recognition and catalysis.

The FAA system from the gut bacterium *Coprococcus eutactus* produces oleoyl aminovaleric acid (OAA), a fatty acid amide that activates several GPCRs expressed in the gastrointestinal tract, including GPR132, GPR119, and PTGER4 (28). The OAA biosynthetic gene cluster encodes three free-standing proteins: OaaA, an adenylation domain that activates oleic acid and transfers it to the ACP OaaACP; and OaaC, a condensation domain that catalyzes amide bond formation between the oleoyl group and aminovaleric acid to produce OAA. Biochemical reconstitution of this pathway demonstrated that OaaC exhibits strict selectivity for small-molecule amine acceptors while tolerating variation in fatty acyl chain length on the donor side (28).

These free-standing NRPS-like proteins may provide structural insights into the catalytic activity and potentially identify features that enable them to function outside of the conventional multidomain NRPS. Here, we present the structural and biochemical characterization of the FAA condensation domain OaaC from *C. eutactus*. Using solution scattering, X-ray crystallography, mass photometry, and in vitro reconstitution assays, we show that OaaC forms an atypical homodimer, both in solution and in the crystal lattice, that appears incompatible with canonical ACP binding modes. We further demonstrate that *holo* and acyl-loaded OaaACP promote dissociation of this homodimer to enable productive catalysis, suggesting a substrate-dependent mechanism for regulating OaaC activity. Finally, phylogenetic and sequence analyses place FAA condensation domains in a distinct evolutionary clade and reveal conserved deviations from the canonical NRPS catalytic motif that may reflect adaptation to small-molecule amine substrates. Together, these results provide new insight into the structure, evolution, and mechanism of FAA NRPS-like condensation domains and establish a framework for understanding how these enzymes function within free-standing biosynthetic systems.

## Results

### OaaC forms a stable homodimer in solution as revealed by SEC-SWAXS

The FAA biosynthetic gene cluster from *C. eutactus* was chosen for initial biochemical and structural analysis based on its unique use of linear small molecule amine and effect on orphan GPCRs. Codon optimized genes encoding OaaA, OaaACP, and OaaC were obtained (Table S1) enabling recombinant expression and purification in *E. coli* (Fig. S1).

To examine the solution state of the free-standing FAA condensation domain OaaC, we performed size-exclusion chromatography coupled to small- and wide-angle X-ray scattering (SEC-SWAXS) (Table S2). OaaC eluted as a single, symmetric peak under all conditions tested (Fig. S2), consistent with a homogeneous species in solution. Scattering profiles collected across the peak exhibited no evidence of aggregation, as indicated by linear Guinier behavior at low *q* and stable radius of gyration values across the elution window (Fig. S2). Calculation of the pair distribution function P(r) yielded an R_g_ of 31.66 Å and a D_max_ of 96 Å (Fig S2), which is larger than that expected for the monomer (R_g_ = 22 Å, D_max_ = 74 Å). Estimation of molecular weight from the SAXS data yielded an apparent mass of approximately 110 kDa, consistent with a homodimeric assembly of OaaC (calculated monomer mass ≈ 54 kDa, dimer ≈ 108kDa). To visualize the architecture of OaaC directly from the raw scattering data, we used DENSS to construct a low-resolution density map *ab initio*. We observed a map (Fig. S2) significantly larger in size than a monomer, and a bilobed shape, consistent with a dimeric architecture (32).

To further assess whether the scattering data were consistent with a monomeric or dimeric architecture, we compared experimental profiles with theoretical scattering curves calculated from multiple structural models using χ² (Fig. 2). Here, χ² represents the reduced chi-squared statistic, which quantifies the agreement between experimental and calculated scattering intensities across all measured *q* values, with lower values indicating better model–data agreement, with an ideal fit resulting in χ² = 1.0. An AlphaFold2 predicted monomeric model of OaaC fit poorly to the experimental data, yielding a χ² value of 878, indicative of substantial deviation between the model and the measured scattering profile (33). In contrast, an AlphaFold2-predicted homodimeric model produced a markedly improved fit (χ² = 5.12), consistent with a dimeric solution state (34). Notably, this dimeric architecture was not guided by prior experimental condensation-domain oligomerization templates, as no previously characterized NRPS condensation domains are known to adopt native homodimeric assemblies.

**Figure 2.**
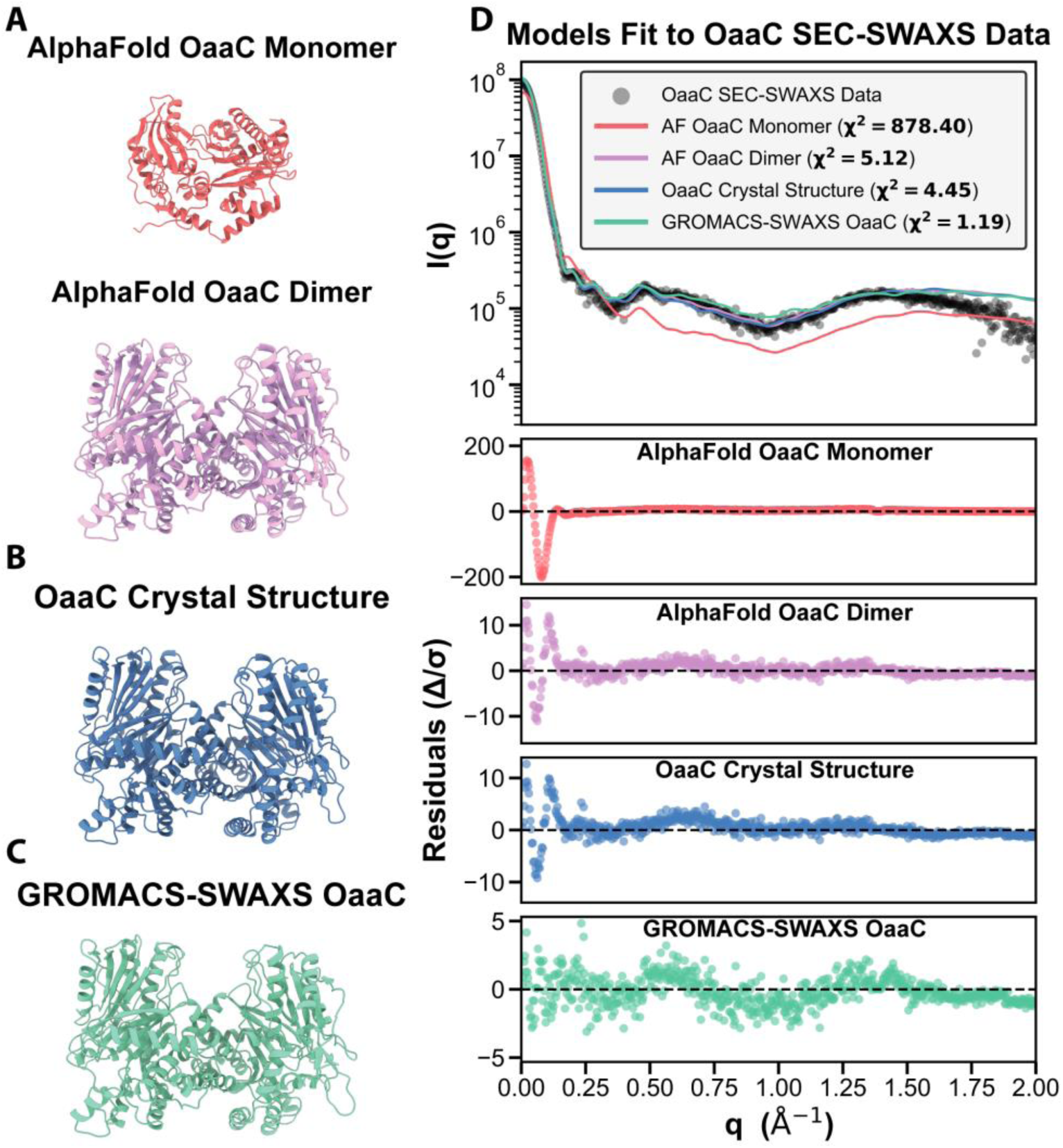
Fitting models to OaaC SEC-SWAXS data. **A,** AlphaFold2 models of OaaC monomer and dimer. **B,** OaaC crystal structure. **C,** GROMACS-SWAXS model of OaaC. **D,** theoretical scattering curves calculated from each structural model fit to experimental SEC-SWAXS data (gray circles). Scattering intensity, I(q) is plotted as a function of the scattering vector q (Å⁻¹) with curves for AlphaFold2 monomer (red), AlphaFold2 dimer (pink), crystal structure (blue), and GROMACS-SWAXS model (teal). χ² values indicate goodness of fit, with lower values representing better agreement with experimental data. Below the I(q) plot are residual plots (Δ/σ) showing the normalized difference between experimental and theoretical scattering for each model.

To further refine the atomistic solution model, we applied GROMACS-SWAXS molecular dynamics refinement, which incorporates the experimental scattering profile as a restraining potential during simulation (35). From the resulting trajectory of refined models, the best-fitting structure yielded a 𝜒^2^ value of 1.19, representing high-quality agreement with the experimental data (Fig. 2C). This was a significantly improved fit over the OaaC AF dimer (𝜒^2^ = 5.12). To determine the structural basis for the improved SWAXS agreement, we compared the representative GROMACS solution model to the OaaC AF dimer. Structural alignments of the isolated monomers demonstrate that while the N-lobe core remains highly stable (RMSD = 0.8Å), subtle inter-domain flexibility results in an overall monomer RMSD of 1.3Å (Fig. S3).

Additionally, the overall complex exhibits a distinct quaternary rigid-body relaxation (global dimer RMSD = 1.91Å). Without altering the fundamental contacts at the interface, the protomers shift slightly in their relative positions, causing the solution dimer to expand relative to the more compact AF model (Fig. S3). Ultimately, this dynamic quaternary breathing allows the complex to achieve excellent agreement with the experimental solution scattering data (𝜒^2^ = 1.19) (Fig. S3C). Together, these SEC-SWAXS analyses indicate that OaaC exists as a stable homodimer in solution under the conditions examined, and that this refined assemble provides the best description of the average solution conformation (Fig. 2).

### Crystal structure of OaaC reveals an atypical homodimer that occludes canonical ACP-binding and donor-access surfaces

We next determined the crystal structure of the free-standing FAA condensation domain OaaC. Crystals diffracted to a resolution of 2.15 Å and belonged to space group H3. The structure was solved by molecular replacement using a single chain of the AlphaFold OaaC model, identifying two OaaC protein chains present in the asymmetric unit (Table S3). Each protomer adopts the canonical condensation-domain fold, consisting of N- and C-terminal lobes arranged in a V-shaped architecture characteristic of NRPS C domains (Fig. 3A). The crystallographic asymmetric unit reveals a homodimeric assembly in which the two protomers associate through an extensive and well-defined interface. The dimeric assembly observed in the experimental crystal structure was consistent with the predicted AlphaFold2 dimeric model.

**Figure 3.**
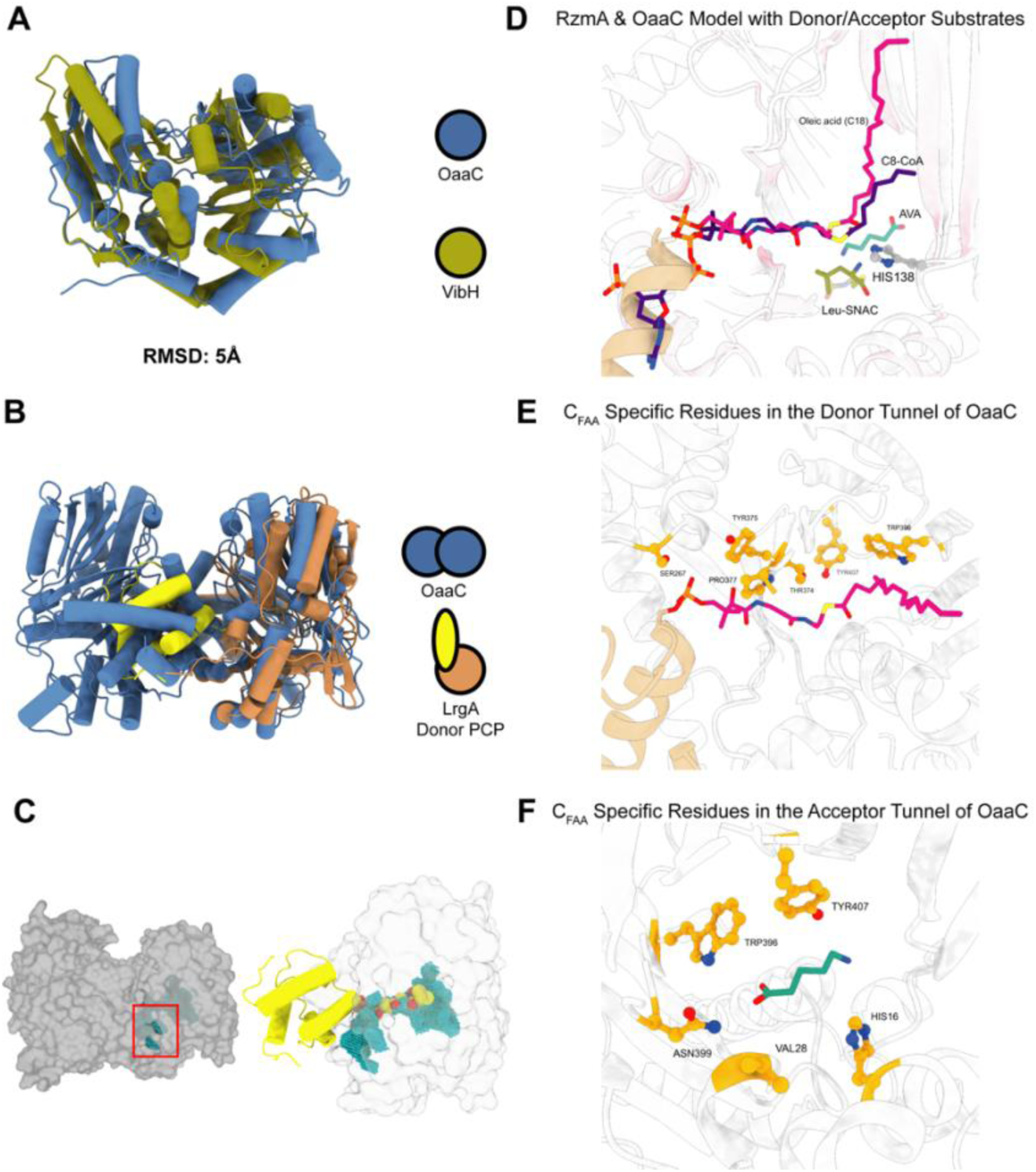
The OaaC homodimer architecture occludes the canonical ACP-binding interface. **A,** Structural superposition of an OaaC protomer with the canonical NRPS condensation domain VibH demonstrates conservation of the overall fold and three-dimensional architecture, consistent with classification of OaaC within the C-domain family. **B,** Overlay of the OaaC homodimer crystal structure with the condensation domain–donor PCP complex from LrgA (PDB ID: 9BE3) reveals that the OaaC dimer interface sterically occludes the canonical donor PCP-binding surface. The spatial arrangement of the second OaaC protomer overlaps with the position occupied by the donor PCP in the LrgA complex, indicating that the crystallographically observed homodimer is incompatible with simultaneous PCP engagement. **C,** Left, surface representation of the OaaC homodimer highlighting partial solvent exposure of the putative donor substrate tunnel (cyan spheres). Right, superposition of the donor PCP from the LrgA complex (yellow ribbons) onto a monomeric OaaC surface model (light gray), illustrating alignment of the predicted donor tunnel (cyan) with the canonical PCP-binding orientation observed in LrgA. **D,** Overlay of a Boltz2 model of OaaC in complex with donor substrate oleic acid loaded-OaaACP and acceptor substrate aminovaleric acid (AVA) with RzmA. The OaaC Boltz2 predicted model with ligands are within proximity of the experimentally solved structure of RzmA with donor and acceptor substrates. **E,** Boltz2 OaaC model’s donor tunnel with the loaded substrate (magenta) and suggestive C_FAA_ specific residues (orange) that are within 4Å of the ligand. **F,** Boltz2 OaaC model donor tunnel with the acceptor substrate (green) and suggestive C_FAA_ specific residues (orange) that are within 4Å of the ligand.

Analysis of the OaaC interface using the Protein Interfaces Surfaces and Assemblies (PISA) server (36) identified this interaction as the most significant interface in the crystal, with a complex formation significance score (CSS) of 1.0. The interface buries approximately 1,400 Å² of solvent-accessible surface area per protomer and involves 42–43 residues from each chain, corresponding to ∼9% of the total residues in each monomer. The interface is composed of a mixture of hydrophobic contacts and polar interactions distributed across both the N- and C-terminal lobes, rather than being restricted to flexible terminal regions, consistent with a specific and stable association. Structural superposition of the two protomers within the dimer reveals minimal conformational differences between chains (root mean square deviation of 0.25 Å over 463 Cα atoms), indicating a largely symmetric assembly (Fig. 3B). The active sites of the two protomers are positioned on opposite faces of the dimer and remain solvent accessible, suggesting that dimerization does not directly occlude the catalytic center itself. Consistent with the SEC-SWAXS analysis, the crystallographic homodimer closely matches the AlphaFold dimer prediction, with comparable protomer orientation and overall dimensions (Fig. 2 B, C). Both models yield similar 𝜒^2^ values compared to the SWAXS data (5.12 vs 4.45), larger than the refined GROMACS-SWAXS model (1.19).

Interestingly, mapping known carrier protein interaction regions onto the OaaC structure reveals that OaaC dimerization sterically obstructs the surfaces that correspond to the canonical donor PCP-binding interface observed in prior structures of complexes of condensation domains with their donor PCP. In the OaaC homodimer, this surface is buried at the dimer interface, rendering it inaccessible for productive ACP docking (Fig. 3B). This configuration contrasts with previously characterized condensation domains, whether free standing or excised from multidomain proteins, that use an interface that appears readily available.

To assess the compatibility of an acyl-ACP interaction with a single chain in the same orientation in canonical NRPSs, we initially modeled this interaction based on known carrier protein binding regions by superimposing OaaC onto the structurally characterized LgrA condensation domain complex. We also generated a predictive model of the ACP-OaaC complex using Boltz2 (37) allowing us to also model the loaded substrate. The Boltz2 prediction yielded an ACP binding interface in excellent agreement with our LgrA-based alignment, choosing the conventional donor site. We evaluated the putative acyl-chain tunnel of OaaC by analyzing the Boltz2 substrate trajectory alongside the superimposed acylated donor ligand from the rhizomide synthetase RzmA (38). This combined analysis reveals a continuous, predominantly hydrophobic tunnel leading from the protein surface to the active site that could readily accept a fatty acyl chain when OaaC is considered as an isolated protomer (38). In the context of the homodimer, however, this donor-access tunnel is partially occluded by the opposing protomer suggesting that dimer formation may restrict the access of ACP-tethered acyl substrates to the catalytic site (Fig. 3B-C), raising the possibility of a dissociation of the dimer for active interaction (see below).

Conventional condensation domains from within multidomain NRPS proteins have been defined to form a “platform” with the neighboring adenylation domain, upon which the loaded carrier protein migrates to deliver substrates and intermediates to the neighboring catalytic domains (5, 39). This consistent arrangement of condensation-adenylation didomains led us to ask if the stable dimeric arrangement of OaaC employed the same face as that observed in multidomain NRPS proteins. Surprisingly, the interface used by OaaC to form a dimer resides on the opposite face of the condensation domain observed in other NRPS condensation-adenylation didomains suggesting that the dimeric conformation observed in OaaC is not simply an alternate approach to bury the same interface on the C-terminal lobe of the condensation domain (Fig. S4). Compared to the condensation domain of the AB3403 structure (40), OaaC contains a 17-residue insertion including a helix from Gln214 to His224 that appears to cover a portion of the condensation domain that would pack against the adenylation domain in a modular NRPS. Together, these observations demonstrate that OaaC forms a stable homodimer through a substantial and energetically favorable interface that preserves the canonical condensation-domain fold but that the observed dimeric conformation appears to sterically interfere with donor ACP access.

### Phosphopantetheinylated OaaACP destabilizes the OaaC homodimer

To directly assess the oligomeric state of OaaC in solution and its response to donor substrates and ACP functional state, we performed mass photometry (Fig. 4). This approach enables label-free measurement of protein mass distributions at nanomolar concentrations and resolves discrete oligomeric species in solution.

**Figure 4.**
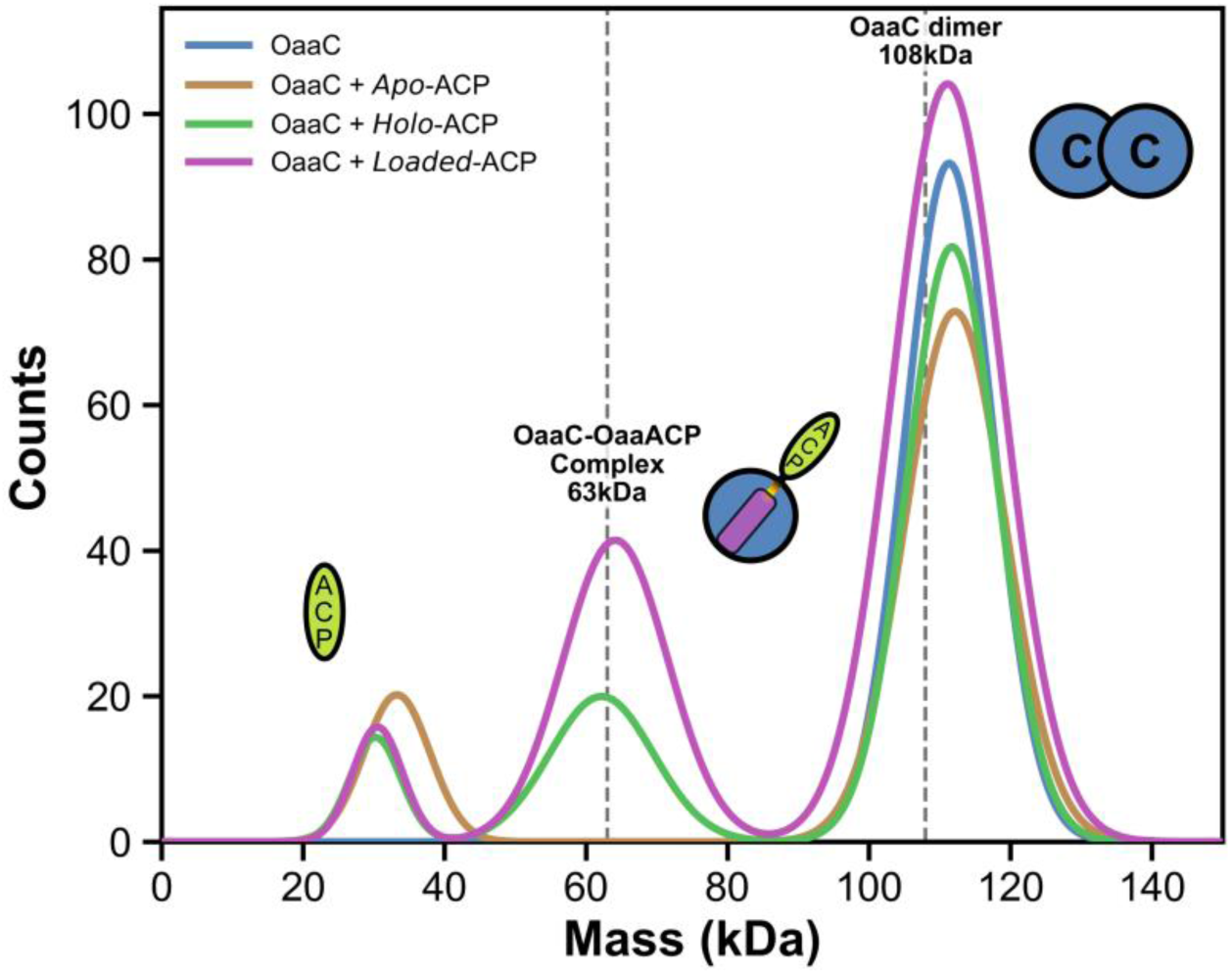
Mass photometry reveals ACP functional state–dependent modulation of OaaC oligomerization and complex formation. Overlayed mass photometry distributions of OaaC alone and in the presence of a ten-fold excess of OaaACP in distinct functional states (*apo*, *holo*, and acyl-*loaded*). OaaC alone exhibits a dominant species corresponding to the homodimer (111kDa ± 6kDa, counts = 847). Addition of *apo*-OaaACP does not measurably alter the oligomeric distribution, with the homodimer remaining the predominant population (112kDa ± 7kDa, counts = 737). In contrast, incubation with *holo*- and acyl-loaded OaaACP results in retention of the homodimer peak but introduces an additional population centered near ∼63 kDa, consistent with formation of an OaaC–OaaACP complex (112kDa ± 6kDa, counts = 725, 62kDa ± 7kDa, counts = 203 and 111kDa ± 8kDa, counts = 1110, 64kDa ± 7kDa, counts = 420 respectively) See Table S4 and Fig. S5 for more details. Reference dashed lines denote theoretical molecular weights for the, OaaC–OaaACP complex (63 kDa), and OaaC homodimer (108 kDa).

Analysis of OaaC alone revealed a single dominant species with a measured mass of 111 ± 6 kDa, consistent with the theoretical molecular weight of the OaaC homodimer (108 kDa). Only a minor population corresponding to monomeric OaaC was detected (Fig. S5), indicating that OaaC exists predominantly as a stable homodimer in solution.

To determine whether free substrates influence OaaC oligomerization independently of ACP, we next examined OaaC in the presence of each substrate individually. Incubation with aminovaleric acid did not alter the mass distribution, which remained dominated by the homodimeric species with no detectable accumulation of monomer (Fig. S5C). Likewise, incubation with oleic acid produced a profile indistinguishable from OaaC alone (Fig. S5D). These results indicate that neither the amine acceptor nor the fatty acid donor substrate destabilizes the OaaC homodimer.

We next evaluated the effect of OaaACP on OaaC oligomerization using a constant ten-fold molar excess of OaaACP over OaaC. Although OaaACP (9.4 kDa) lies below the reliable detection limit of the 2MP instrument, independent measurements of apo-, holo-, and acyl-loaded OaaACP reproducibly yielded weak signals in the 29–35 kDa range (Fig. S5E-F). These low-intensity features fall below the instrument’s quantitative detection threshold and were not considered in subsequent interpretation.

Addition of *apo*-OaaACP did not measurably alter the mass distribution of OaaC, which remained dominated by the homodimer (Fig. 4). In contrast, incubation with *holo*-OaaACP resulted in the emergence of a distinct intermediate-mass species at 62 ± 7 kDa, consistent with formation of an OaaC–OaaACP complex. A similar distribution was observed upon incubation with acyl-loaded OaaACP, obtained by loading *apo*-OaaACP with oleoyl-CoA and the promiscuous *sfp* phosphopantetheinyl tranferase to avoid the need to include OaaA and the potential for incomplete loading (Fig. 4). Incubation of OaaC with loaded OaaACP resulted in a prominent species at 64 ± 7 kDa alongside the homodimer peak. The measured intermediate masses are consistent with the expected molecular weight of the OaaC–OaaACP heterocomplex (∼63 kDa).

*Holo*- and acyl-*loaded* OaaACP produced a stable species with a mass corresponding to the OaaC–OaaACP complex. In contrast, apo-OaaACP and free substrates failed to induce detectable complex formation. The reproducible emergence of this intermediate-mass population under phosphopantetheinylated ACP conditions indicates that ACP modification state governs engagement with OaaC and promotes dissociation of the homodimer. Notably, this functional behavior is consistent with the structural observation that homodimerization occludes the canonical ACP-binding surface, suggesting that ACP engagement requires disruption or remodeling of the dimer interface, providing a potential modification state and regulation of condensation-domain oligomerization.

### LC–MS reconstitution demonstrates OaaC-catalyzed fatty acid amide formation across multiple acyl-chain lengths

We next reconstituted the pathway using purified OaaC, OaaA, and OaaACP in the presence of fatty acid and amine substrates and analyzed reaction products by LC-MS (Fig. 5A). Reactions with individual combinations of fatty acyl donors and amine acceptors were incubated for 2 h prior to analysis. LC-MS analysis of reconstituted reactions revealed the formation of discrete products with retention times and extracted-ion masses consistent with fatty acid amides derived from the supplied substrates. Overlaid extracted-ion chromatograms (EICs) showed robust formation of oleoyl-, palmitoleoyl-, and lauroyl-linked amides when aminovaleric acid or γ-aminobutyric acid was provided as the amine acceptor (Fig. 5B). The detection of products derived from fatty acids differing in chain length and degree of unsaturation indicates that both OaaA and OaaC can accommodate a range of acyl-chain substrates under the assay conditions. Each FAA product exhibited a distinct retention time corresponding to its acyl-chain composition, and no comparable peaks were detected in control reactions lacking OaaC (Fig. S7), indicating that product formation is dependent on the condensation domain.

**Figure 5.**
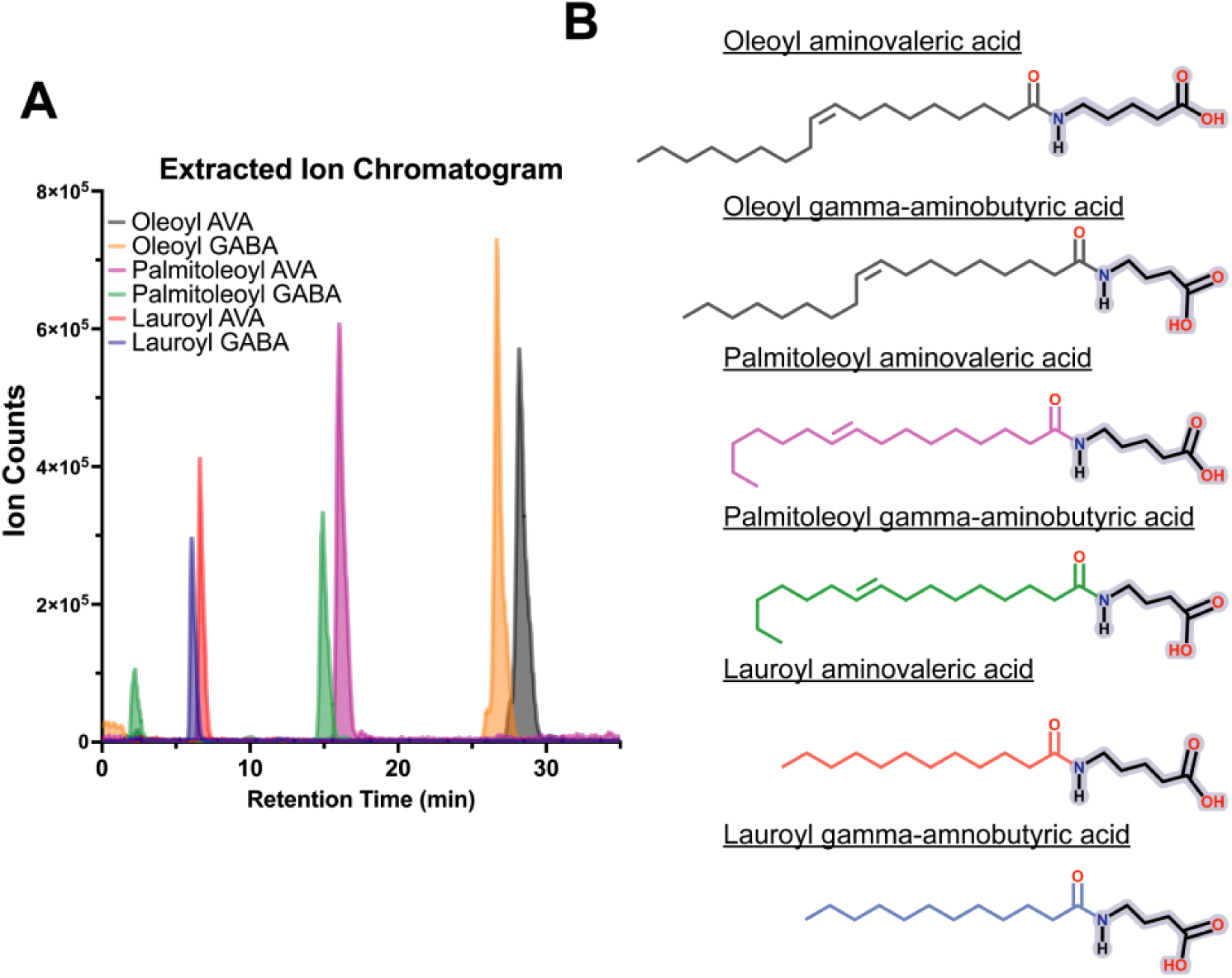
LC-MS analysis of OaaC-catalyzed fatty acid amide formation. **A,** Schematic of the in vitro reconstitution of the fatty acid amide (FAA) pathway, in which OaaC, OaaA, and OaaACP are combined with fatty acid and amine substrates and analyzed by LC-MS. **A,** Overlaid extracted-ion chromatograms (EICs) from individual reactions, showing the formation of distinct FAA products. **B,** Chemical structures of the expected FAA products following a 2-hour incubation. Acyl-chain coloring corresponds to the associated EIC traces shown in (4B).

Together, these LC–MS data demonstrate that OaaC catalyzes fatty acid amide bond formation *in vitro* and is capable of utilizing fatty acyl substrates spanning multiple chain lengths and saturation states (Fig. 5). These results establish a biochemical foundation for linking the substrate-dependent modulation of OaaC oligomerization observed in structural and biophysical experiments to catalytic activity.

### Phylogenetic analysis places FAA condensation domains in a distinct clade related to C_Starter_ domains

To evaluate how FAA-associated condensation domains relate to canonical NRPS C-domain families, we used the Enzyme Function Initiative Enzyme Similarity Tool (EFI-EST) (41) to assemble a sequence set containing curated representatives of major C-domain functional classes together with 127 FAA-associated C domains identified. We combined a selection of FAA C domains with a dataset of diverse condensation domains described previously (42). Maximum-likelihood phylogenetic reconstruction was performed using IQ-TREE (43), and trees were visualized with iTOL (44). This analysis revealed that FAA-associated C domains form a well-supported monophyletic clade that is distinct from canonical NRPS C-domain classes (Fig. 6A). Within the broader condensation-domain phylogeny, the FAA clade is most closely related to C_Starter_ domains (Fig. 6A), consistent with a shared feature of utilizing an acyl-chain donor substrate. Notably, FAA systems differ from C_Starter_ domains in both donor-carrier chemistry and acceptor substrate class. FAA pathways utilize an acyl-loaded ACP donor and condense acyl chains with freely diffusing small-molecule amines, whereas C_Starter_ domains typically accept acyl-CoA donors and act on aminoacyl acceptors presented by a downstream carrier protein, showing that FAA condensation domains form a distinct evolutionary subgroup within the broader NRPS condensation-domain superfamily (Fig. 6A).

**Figure 6.**
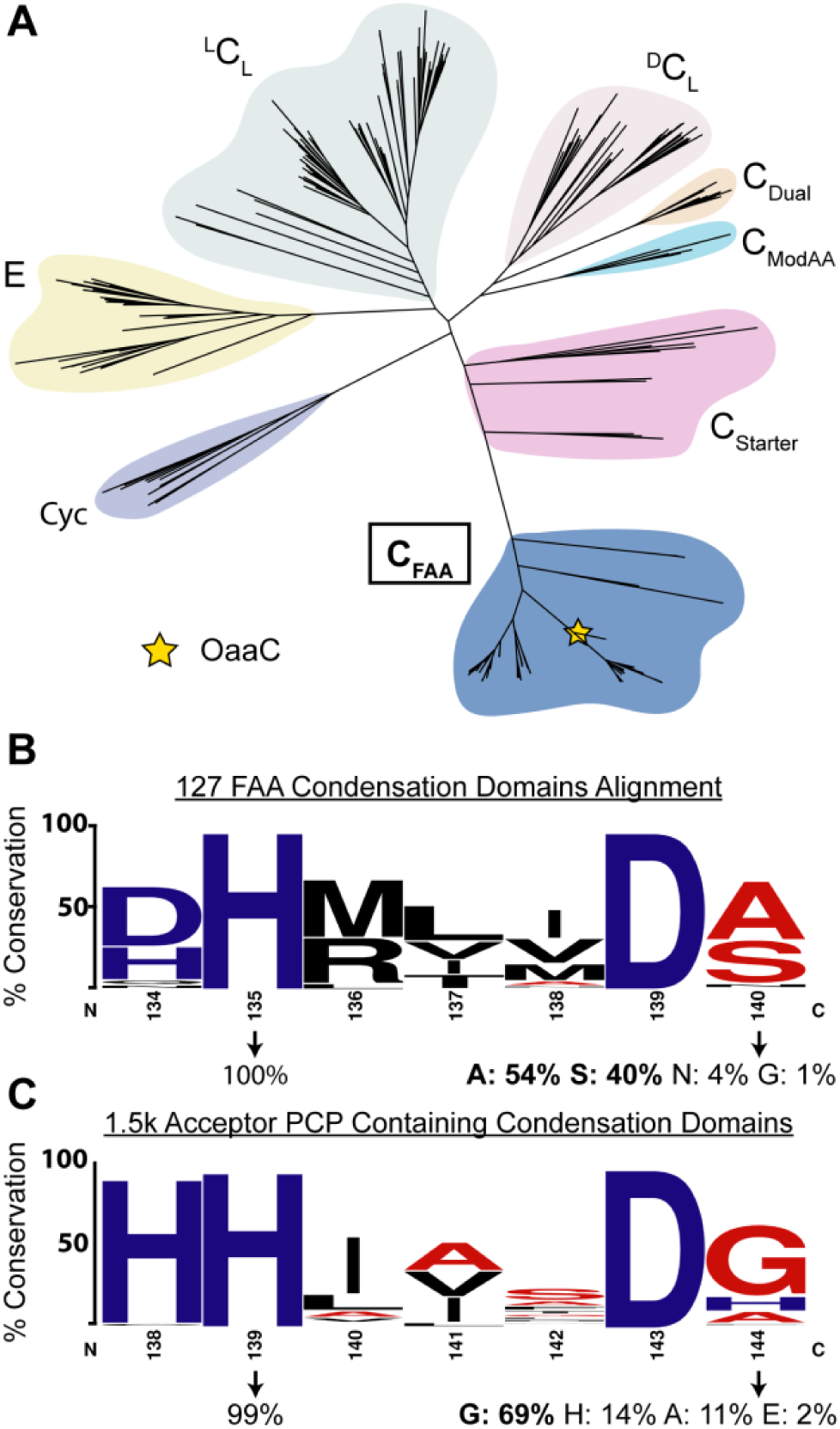
Phylogenetic and motif analyses of FAA condensation domains. **A,** Maximum-likelihood phylogeny of representative NRPS condensation (C) domain classes together with 127 FAA-associated C domains. FAA C domains form a distinct monophyletic group, with C_Starter_ domains as their nearest neighboring clade. OaaC is indicated with a yellow star. **B**, Sequence logo of 127 FAA C domains showing conservation of the first histidine and variability at the fourth motif position, which is enriched for serine and alanine. **C**, Sequence logo of PCP-dependent condensation domains highlighting predominant glycine usage at the fourth motif position

To determine whether sequence-level features distinguish FAA condensation domains from canonical NRPS C domains, we analyzed conservation of the catalytic motif across the 127 FAA-associated sequences used in the phylogenetic reconstruction in comparison to 1500 canonical C-domain sequences obtained from InterPro (45) that were selected to ensure they contained upstream and downstream PCP domains. C_FAA_ domains like OaaC contains a (D/H)HxxxDS motif that differs from the canonical HHxxxDG motif commonly associated with PCP-dependent NRPS condensation domains. Multiple sequence alignment of the FAA set shows that the second histidine position is highly conserved (>99%) as is the aspartic acid at the sixth position. In contrast, the residue following the aspartic acid displays substantial variability (Fig. 6B). In the FAA sequence set, this position is most frequently occupied by serine (42%) or alanine (36%), while the glycine common in canonical NRPS condensation domain occurs in only 17% of sequences (Fig. 6B).

We next attempted to see if C_FAA_ possessed distinguishing features relative to ^L^C_L_ domains. We reasoned that C_FAA_ domains utilizing a small molecule amine and fatty acid may have some sequence/structural features that are emphasized when compared to ^L^C_L_ domains. Using MiBIG we gathered annotated sequences from the major subtypes of condensation domains (^L^C_L,_ ^D^C_L,_ C_Starter,_ C_Glyc,_ C_Dual,_) and added C_FAA_ to this subtype list (46). Then by performing subtype specific sequence alignments we could observe consensus residues per subtype. C_FAA_ and ^L^C_L_ consensus sequences were aligned and mapped to residue distributions at each position. We defined a threshold for potential C_FAA_-specific residues that were highly conserved (>85%) but occurred infrequently (<25%) in the ^L^C_L_ subtype. Using a Boltz2 model of OaaC with donor and acceptor substrate bound, we then looked at residues that met this threshold and were also present in putative donor and acceptor sites (Fig. 3E, 3F). This subtype sequence to structure analysis suggests that C_FAA_ domains have conservation of some hydrophobic residues that line the putative donor tunnel (Fig. 3E). Similarly, the acceptor site contains five residues that are suggested to be highly conserved and predominantly present in C_FAA_ domains relative to ^L^C_L_ domains (Fig. 3F). These bioinformatic analyses suggest that OaaC and C_FAA_ domains broadly may have made specific changes not only at their catalytic motif but also in their donor and acceptor tunnels. This agrees with their divergence from other subtypes in our phylogenetic analysis and being most closely related to C_Starter_ domains that typically utilize acyl-CoAs.

## Discussion

Condensation domains of NRPSs catalyze amide bond formation through the attack of the amine of the acceptor substrate on the thioester bond of the donor substrates tethered to upstream carrier proteins. Although the catalytic HHxxxDG motif and overall fold are highly conserved, increasing structural and biochemical evidence indicates that C domains exhibit substantial diversity that extends beyond canonical peptide bond formation (47, 48). The present study provides structural, biochemical, and biophysical characterization of OaaC, a free-standing condensation domain involved in the biosynthesis of fatty acid amides (FAAs) in gut bacteria. These findings expand the functional landscape of the NRPS C-domain superfamily and provide insight into how compact NRPS-like pathways generate lipid-derived signaling molecules.

We determined that OaaC adopts a stable homodimeric architecture that sterically occludes the canonical carrier-protein binding interface. Structural comparisons with previously characterized NRPS condensation domains indicate that the overall fold of OaaC is conserved, but the quaternary organization is incompatible with conventional donor carrier proteins. Our mass photometry data support a dynamic equilibrium between dimeric and heterodimeric complexes in solution influenced by the functional state of OaaACP. Specifically, *holo*- and acyl-loaded OaaACP promote the formation the OaaC-OaaACP heterodimer consistent with the productive enzyme-carrier protein interaction. These observations support a model in which OaaC adopts an inaccessible dimeric state that is disassociated once *holo* or loaded carrier protein interacts (Fig. 7). Such a mechanism would potentially suppress unproductive interactions with other *apo* carrier protein or CoA thioesters while coupling catalysis to the presence of the specific loaded donor substrate.

**Figure 7.**
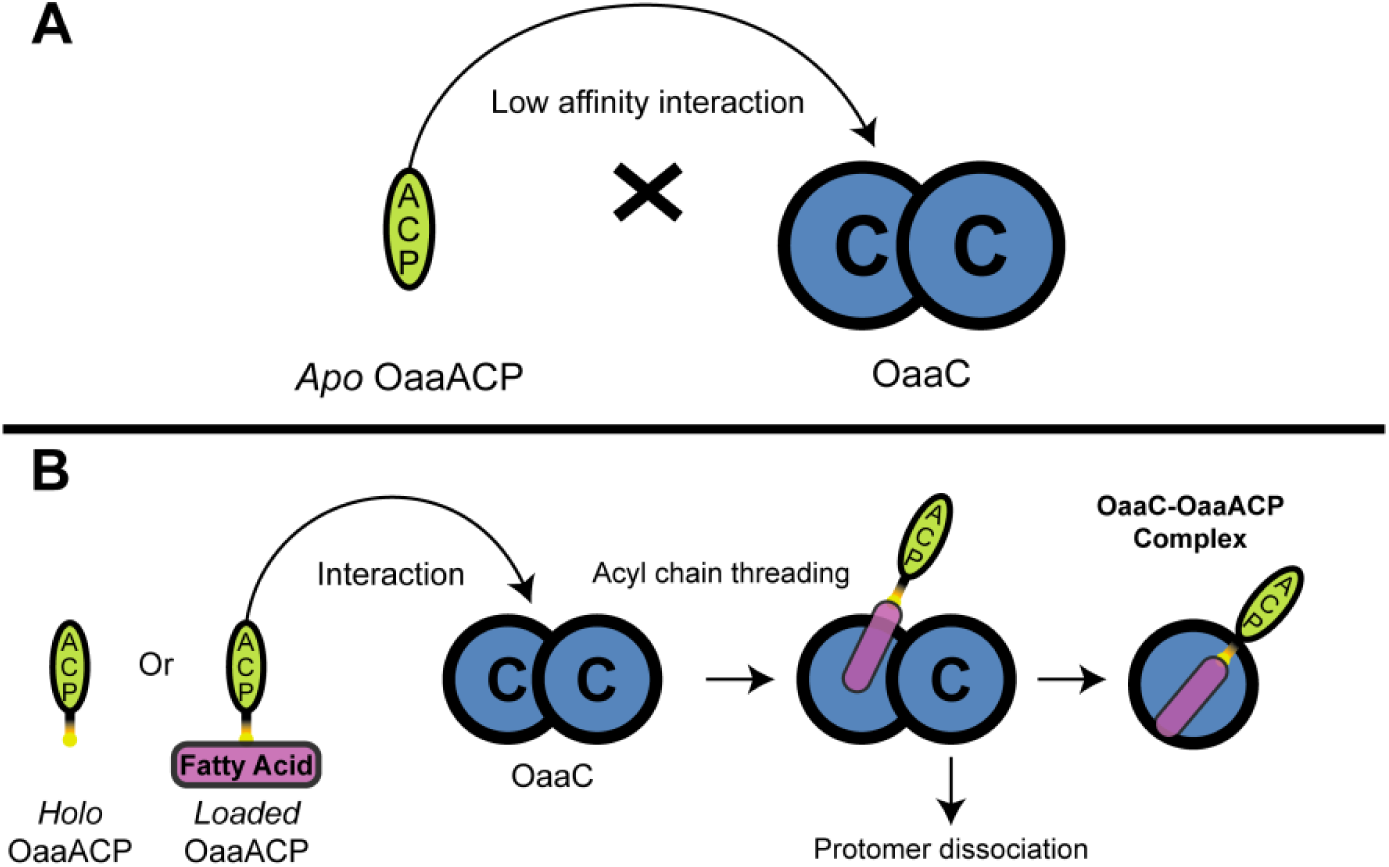
Proposed model for autoinhibitory dimerization and activation of an FAA condensation domain. **A,** OaaC forms an autoinhibitory homodimer in which the *apo*-OaaACP interacts weakly and is unable to productively dock onto either condensation protomer. **B,** In contrast, *holo* or acyl-*loaded* OaaACP engages the OaaC homodimer, enabling threading of the acyl chain into one protomer. This interaction is proposed to promote dissociation of the homodimer, allowing formation of a OaaC–OaaACP complex at the donor-site interface.

This model aligns with emerging evidence that condensation domains can be subject to regulatory mechanisms that extended beyond the catalytic motif. Structural and mechanistic studies have demonstrated that C domains impose stringent donor and acceptor substrate selectivity and frequently represent bottlenecks in engineered NRPS systems (48, 49). Recent work on lipoinitiating and starter condensation domains has shown that acyl-chain selectivity can be tuned through mutations within the donor binding tunnel, highlighting the role of structural features outside the catalytic motif in determining substrate specificity (38, 50). The structural organization observed for OaaC suggests that oligomerization may represent an additional layer of control in NRPS-like enzymology. For free-standing FAA NRPS enzymes that operate outside of the canonical multidomain assembly-line architecture, conformational gating through oligomerization may provide a mechanism to maintain specificity and catalytic efficiency in the absence of enforced domain proximity.

Our phylogenetic analysis of OaaC supports the concept that FAA-associated condensation domains represent a specialized branch of broader C-domain superfamily. Phylogenetic analysis places these enzymes near starter condensation domains, which catalyze acyl-transfer reactions during lipopeptide biosynthesis (Fig. 6). This relationship is intuitive, mechanistically speaking, as both classes utilize fatty-acyl donor substrates. However, FAA associated condensation domains have specificity for freely diffusing amines rather than carrier protein bound amino acids. The sequence divergence and altered active site architecture observed in these enzymes likely reflect an adaptation to this unusual donor acceptor pairing (Fig. 6B). We note, also, that initiating condensation domains that install a salicylate or 2,3-dihydroxybenzoic acid moiety that are common in many peptide siderophores also cluster with these starter condensation domains, indicating that this organization does not simply result from the fatty acyl substrate of the FAA and lipoinitiating condensation domains (51, 52). These findings support the variety of condensation domains chemistries and that diversification of this enzyme family underpins much of the chemical diversity in NRPS derived natural products (47).

Beyond enzymology, these findings have implications for understanding the biosynthesis of signaling molecules produced by the human gut microbiota. Several gut bacterial species encode compact NRPS-like pathways that generate *N*-acyl amides capable of interacting with mammalian G-protein-coupled receptors (GPCRs) and influencing host physiological processes (28). These microbial metabolites can mimic endogenous lipid signaling molecules and activate host receptors involved in immune regulation, metabolism, and intestinal homeostasis (53, 54). The biochemical behavior of OaaC suggests that FAA production may be tightly coupled to the metabolic state of the producing organism. Given the broad specificity of the adenylation domain that we observe and in agreement with prior studies (28), it remains to be seen if different loaded OaaACP proteins show differing capacities to trigger the formation of competent OaaACP-OaaC complexes. Additionally, we did not observe product formation in reactions of Oleoyl-CoA with OaaC, suggesting that the combined activities of the adenylation and condensation domains are necessary to increase selectivity of the system that would not be observed if OaaC could directly load the fatty acid from an acyl-CoA onto aminovaleric acid (Fig. S9). Because productive interaction between OaaC and OaaACP appears to depend on carrier-protein functionalization and acyl loading, FAA biosynthesis may be regulated by intracellular fatty-acid availability and phosphopantetheinylation status. Such coupling would enable bacteria to modulate production of host-active metabolites in response to nutrient availability or metabolic conditions within the gut environment.

More broadly, mechanistic insight into FAA biosynthesis provides an opportunity to better understand how microbial secondary metabolism interacts with mammalian signaling pathways. Microbiome derived small molecules are increasingly recognized as important mediators of host physiology, yet the structural and enzymatic mechanisms that control their production remain poorly understood (53). Structural characterization of enzymes such as OaaC therefore provides a mechanistic explanation for how metabolic flux dictates production of host-interacting metabolites. Specifically, the transition between an autoinhibited dimer and a functional heterodimer suggests that FAA production is gated by the availability of both holo-and acyl-loaded carrier proteins, ensuring these signaling molecules are only synthesized when the microbial metabolic state and biosynthetic machinery can support it.

From an engineering perspective, the features identified in the FAA C domain OaaC may also inform efforts to reprogram NRPS-derived biosynthetic systems. Currently, much of the emphasis is placed on the adenylation domain, with many studies being able to successfully engineer the “gatekeeping” domain only to be met with low total product yields (48, 55). Other engineering methods focus on C-domain swapping, module swapping, or altering the substrate specificity of the C domain (15, 38, 49, 50). However, the structural organization of OaaC demonstrates that productive catalysis is governed not only by substrate specificity but also by quaternary structural dynamics that regulate carrier-protein engagement. Our results support the need for continued study to optimize carrier-protein recognition and structural gating mechanisms. The minimal architecture of FAA biosynthetic systems may therefore represent an attractive platform for engineering novel amide-linked metabolites or lipid-modified molecules.

Future studies will be necessary to further define the mechanistic basis of OaaC regulation. Mutational analysis of residues within the dimer interface could determine whether disruption of oligomerization alters catalytic activity or carrier-protein interaction and, potentially, plays a role in acylamide formation in the cell. Structural or solution-based approaches such as cryo-EM, SWAXS, or cross-linking mass spectrometry may help define the architecture of the OaaC–OaaACP complex and clarify how carrier-protein engagement influences enzyme conformation. In addition, systematic variation of acyl donor and amine acceptor substrates could further define the determinants of substrate specificity in FAA-associated condensation domains. Together, such studies will provide a more complete understanding of how these enzymes control the production of microbiome-derived signaling molecules and may enable rational engineering of FAA biosynthetic pathways.

In summary, the present study establishes OaaC as a structurally and mechanistically distinct member of the NRPS condensation-domain superfamily and provides a structural framework for understanding fatty acid amide biosynthesis in gut bacteria. The observed oligomeric organization suggests a potential regulatory mechanism that couples catalytic activity to carrier-protein functionalization, thereby linking metabolite production to intracellular metabolic state. These findings expand the mechanistic diversity of condensation domains and highlight the role of NRPS-like enzymes in the generation of microbiome-derived signaling molecules.

## Methods

### Protein expression

Codon optimized genes encoding OaaA, OaaC, OaaACP inserted into a pET28b_HisTEV plasmids containing a kanamycin resistance marker and a N-terminal 5xHis-tag were purchased from Twist Bioscience (Table S1, Table S5). Plasmids were transformed into NEB turbo cells for storage and into *E. coli* BL21-BAP1 cell line. Large scale expression of each enzyme was done by inoculating 1L of LB medium with kanamycin (50 µg/ml) with 1% inoculum from a primary culture. Cells were grown at 37°C and 250 rpm and monitored until the optical density reached 0.5; flasks were placed in the cold room (4°C) for 20 minutes. After cooling, isopropyl-𝛽-D-thiogalactopyranoside (IPTG) was added at a final concentration of 0.5 mM and cells were grown at 20°C, 200rpm for a total duration of 18-20 h. Cells were harvested by centrifugation and stored at -80°C for further experiments.

### Protein purification

Cell pellets were resuspended in 10 mL lysis buffer (50 mM HEPES pH 7.5, 150 mM NaCl, 20 mM imidazole, 0.2 mM TCEP, 3% glycerol) per g of pellet. The dissolved pellet was then sonicated in 50-60 mL aliquots using the following protocol: 7 seconds on, 10 seconds off, 60% amplitude, and total time of 15 minutes. Lysate was then ultracentrifuged at 185,677 × g at 4°C for 40 minutes. The supernatant was extracted and filtered through a 45 µ filter into a clean beaker.

Sample was then passed over a 5 mL nickel affinity His-trap column, using an BioRad FPLC at 2 ml/min and eluted with elution buffer (50 mM HEPES pH 7.5, 150 mM NaCl, 300 mM imidazole, 0.2 mM TCEP, 3% glycerol) at 4 mL/min. The sample was then placed in an appropriate molecular weight cut off dialysis bag along with TEV protease (4mg/mL) to cleave the histidine tag. This bag was placed in dialysis buffer (50 mM HEPES pH 7.5, 150 mM NaCl, 20 mM imidazole, 0.2 mM TCEP, 3 % glycerol) at a 1:10 sample to dialysis buffer ratio for 18-20 hours. Dialyzed sample was then put back through the IMAC column and the flow-through was collected and concentrated for loading onto a gel filtration column. A HiLoad 16/600 superdex 200 pg or a HiPrep 26/60 Sephacryl S-300 HR column were used for each proteins final gel filtration purification step. All proteins were eluted at 1-1.5 mL/min using a gel filtration buffer (10mM HEPES pH 7.5, 50mM NaCl, 0.2mM TCEP) and flash frozen in liquid nitrogen and stored at -80°C.

### Liquid chromatography mass spectrometry (LC-MS): Product formation Assay

Product formation reactions were performed in 100 µl and contained 25 mM MgCl_2_, 5 mM ATP, 1 mM TCEP, 1 mM amine, 1 mM fatty acid, 1 mM CoA, 1 µM sfp, and 10 µM each of OaaA, OaaC, OaaACP. The reaction was incubated in a shaker at 250 rpm and 23°C for 2 h. Reactions were quenched with an equal volume of acetonitrile. Samples were then centrifuged for 10 minutes at 14,000 rpm in a microfuge and allowed to sit at room temperature for 10 minutes. 5uL of the supernatant was injected onto an Agilent 1200 Infinity II with an attached single quad mass spectrometer using a C18 4 µm reverse phase column (4.6x100mm). An isocratic elution using 20% HPLC water (A) and 80% acetonitrile (B) was used at a flow rate of 0.5 mL/min to separate the fatty acid amide product. Extracted ion chromatograms were examined for expected products.

### SWAXS data collection and processing

OaaC samples were sent to the NSLS-2 LiX beamline for SEC-SWAXS data collection. Sample conditions are described in Table S2. The raw image data was auto processed with the LiX data processing pipeline (56). Radially averaged profiles were obtained and then manually buffer subtracted using the BioXTAS RAW software suite (57). The buffer subtracted profiles were then subjected to Guinier analysis and molecular weight calculation via volume of correlation and Porod volume in RAW (57, 58). Indirect Fourier transformation was used to calculate the P(r) curve using the denss-fit-data script (59). Data collection statistics are presented in Table S2. Twenty *ab initio* density maps were calculated, aligned and averaged with DENSS v1.8.6 (32, 60). Density maps are visualized as volumes using PyMOL (61). The denss-pdb2mrc script was used to calculate profiles from atomic models and perform fitting to OaaC SWAXS data (32, 62).

### OaaC crystallization

OaaC was subjected to multiple crystallization screens at a protein concentration of 13 mg/mL. Crystals were observed in several conditions from the MCSG-1 (Hampton Research) crystallization screen. The most promising crystals were observed from drops containing 0.1 M CHES, NaOH, pH 9.5, 30% w/v PEG 3000. This condition was optimized using a 24-well hanging drop plate with OaaC at 12 mg/mL and a crystallization cocktail of 0.1 M CHES, NaOH, pH 9, 3mM AVA, 35% PEG 3000 w/v, and a protein:cocktail drop volume ration of 2:1. Crystals were harvested and soaked in fresh mother liquor containing 3 mM AVA for 20 seconds.

### Crystal structure Solution

X-ray diffraction data was collected remotely at the SSRL and APS beamline on multiple crystals. Data processing was performed using the Phenix package (63). The OaaC ligand-soaked dataset was processed to a resolution of 2.15Å in a space group of H3. Structure solution was performed by using molecular replacement in which an AlphaFold2 model of a single protein chain was used as the initial search model for the Phaser module in Phenix(33). The structure identified two protein chains in the asymmetric unit. Model building was performed using Coot and subsequent refinements were done using phenix.refine (64). Although some weak density was visible in the active site, it was insufficient to model the soaked ligand.

### GROMACS-SWAXS simulation

GROMACS-SWAXS simulations were performed using an AlphaFold model as the starting structure and the experimental SWAXS data of OaaC as the target profile (35). The protein was placed in a 10x10x10 nm cubic simulation box, solvated, and energy-minimized for a maximum of 5,000,000 steps (or until a force tolerance of 1000 kJ/mol/nm was reached).

Following minimization, the system was equilibrated for 500 picoseconds. The SAXS-driven production simulation was then run for a total of 2 nanoseconds, yielding a profile that fit the experimental data with a χ² of 1.19 (initial χ² was 14.6).

### Bioinformatic analysis

To establish the evolutionary context of FAA condensation domains, we constructed a sequence similarity network (SSN) and performed a phylogenetic analysis. Using the Enzyme Function Initiative tools (41), we built an SSN seeded with OaaC. This SSN, which consisted of speculative FAA condensation domains, was then filtered by length to only include sequences that were within the 300-520 amino acid range. This resulted in ∼300 sequences which also captured other known FAA C domains. Analysis of neighboring genome neighborhood regions with EFI showed that the identified protein sequences contained free-standing adenylation and carrier protein domains. A multiple sequence alignment was carried out using the Muscle alignment server (65), and sequence alignments were visualized using Jalview and WebLogo. To construct the database consisting of C domains that use an acceptor PCP, we use the Interpro database to locate ∼1800 NRPS clusters that had an architecture which included an acceptor PCP (45). The Muscle server was then used again to align the acceptor-containing NRPS proteins, and the resulting alignment was visualized using the software stated above. The phylogenetic analysis was performed using a data set of condensation domains described previously (42) into which we added over 100 FAA condensation domain sequences identified through EFI analysis. The resulting phylogenetic tree was visualized with the interactive Tree of Life (iTOL) visualization tool (44).

To build a robust dataset for comparing C_FAA_ and conventional NRPS Condensation (C) domains, protein sequences were systematically retrieved from public databases and strictly filtered. C domains were extracted from the MIBiG database (v4.0). Domain boundaries and preliminary subtype classifications (e.g., ^L^C_L_, ^D^C_L_, C_Dual_, C_Starter_, C_Glyc_) were parsed from GenBank annotations (46). To ensure structural homology and domain integrity, parent protein sequences were queried against the PF00668 Hidden Markov Model (HMM) using hmmsearch (66) Sequences were retained for downstream analysis only if they exhibited a minimum HMM model coverage of 70% and a high degree of overlap (≥ 90%) with the annotated MIBiG boundaries. External, manually curated FAA-like C-domains were appended to this dataset.

To identify the stable, defining structural core of each domain class, filtered sequences were profiled by subtype to extract un-gapped consensus architectures. Filtered sequences were segregated by subtype and subjected to Multiple Sequence Alignment (MSA) using MAFFT (67) To isolate the stable structural core of each subtype and discard highly divergent loop regions, a strict trimming protocol was applied. Any alignment column with less than 70% non-gap occupancy was dropped. The most frequent amino acid in each of the remaining columns was extracted to generate an un-gapped, representative core consensus sequence for each subtype.

To pinpoint the specific evolutionary divergences that dictate substrate specificity, core consensus sequences were globally aligned and statistically evaluated to identify Specificity Determining Positions (SDPs). To establish a universal spatial grid for comparative analysis, the ungapped core consensus sequences of all subtypes were globally aligned using MAFFT (67).

This shared alignment grid allowed for direct pairwise comparisons between the FAA architecture and comparison subtypes (e.g., ^L^C_L_). At each shared position, the evolutionary relationship was classified based on the chemical properties of the top consensus residues. Positions were designated as "highly conserved" if they maintained an intra-subtype conservation of ≥ 85% and a non-gap occupancy of ≥ 80%. To identify structural features unique to the FAA architecture, a distribution filtering strategy was employed. A residue was defined as an FAA-specific structural determinant if it met two strict criteria: **1** the residue was highly conserved (≥ 85%) within the FAA subtype, and **2** the identical amino acid appeared in ≤ 25% of the entire sequence distribution of the comparison subtype. This ensured that highlighted determinants were genuinely rare architectural divergences rather than simple variations in top consensus preferences.

To physically contextualize these sequence-based determinants within a three-dimensional space, the identified SDPs and conserved landmarks were mapped onto OaaC structural models. The FAA core consensus sequence was pairwise aligned to the sequence of the OaaC structural models (standalone and ligand-bound complexes) to translate sequence coordinates into 3D spatial coordinates. Conserved core landmarks and identified SDPs were programmatically mapped onto the OaaC structures. For the complex models, residues were further filtered by their proximity to bound ligands (LIG1, DAV/LIG2), with contacts defined by a ≤ 4 Å distance cutoff. All structural visualizations and mapping overlays were generated autonomously using Python-generated ChimeraX command (.cxc) scripts.

### Mass photometry

Mass photometry was performed with a Refeyn 2MP instrument. The system was allowed to come to operating temperature and balance for a minimum of two hours. Sample slides were cleaned with isopropyl alcohol and rinsed twice with MQ before placing the sample cassette on the slide. Using a 5x stock of the MFP1 calibrant we added 10uL buffer and 10uL of the calibrant and collected our protein standard measurement always within 1-1.5 hours of sample measurements. For experimental samples, we used 15 µL of buffer to blank the slide followed by 5 µL of sample, a 7s pause after lid closure, then the measurement was taken. To capture the OaaC-OaaACP complex, a sample at 1:10 ratio at µM concentration was first preincubated for 90 minutes at room temperature. Then followed sequential 1:100 and 1:10 dilutions in a 1.5mL Eppendorf tube. The final diluted sample was used for all subsequent measurements at a final concentration of OaaC and OaaACP of 2.5nM and 25nM, respectively. The data were analyzed initially using the Refeyn Acquire software then saved and replotted using the Matplotlib library in python3 for visualization in this manuscript.

## Data Availability

The crystal structure of OaaC has been deposited with the worldwide Protein Data Bank with PDB ID: **12KH.**

SASBDB ID: Submitted, awaiting ID assignment.

## Supporting Information

Supporting information is available at *website to enter*.

## Acknowledgements

We thank Dr. Shirish Chandokar for assistance with data collection at the LiX beamline 16-ID at NSLS-II. We would like to thank Patrick K. Oduro for his guidance on running GROMACS-SWAXS simulations. Additionally, we thank Dr. Ketan K. Patel for his many insightful conversations and training in protein expression and purification.

## Authorship Contributions

The experimental work described in the manuscript was performed by J.S under the supervision of A.M.G and T.D.G. Data analysis and interpretation were performed by J.S, A.M.G and T.D.G. All authors wrote, edited, and revised the manuscript.

