## Supplementary Information for "Structure of a Stand-Alone Homodimeric NRPS Condensation Domain Reveals Occlusion of the Canonical Carrier-Protein Interface"

### Supporting Information

**Table of Contents**

Figure S1. SDS-Gel of Purified Proteins. .... S2
Figure S2. Analysis of OaaC SEC-SWAXS Data .... S3
Figure S3. Structural basis of OaaC solution-state relaxation .... S4
Figure S4. Comparison of OaaC dimer with an NRPS module .... S5
Figure S5. Mass Photometry .... S6
Figure S6. Mass Photometry Histogram Distribution .... S7
Figure S7. LC-MS production of oleoyl aminovaleric acid. .... S8
Figure S8. Oleoyl aminovaleric acid production with Sfp loaded OaaACP .... S9
Figure S9. Assay of OaaC with fatty acyl CoA. OaaC is unable to use Oleoyl CoA .... S10
Supplemental Tables .... S11
Table S1. Codon optimized genes used. .... S11
Table S2. SAS sample details .... S12
Table S3. Data collection and refinement statistics .... S14
Table S4. Mass photometry statistics .... S15
Table S5. Protein sequences .... S16

### Supplemental Figures

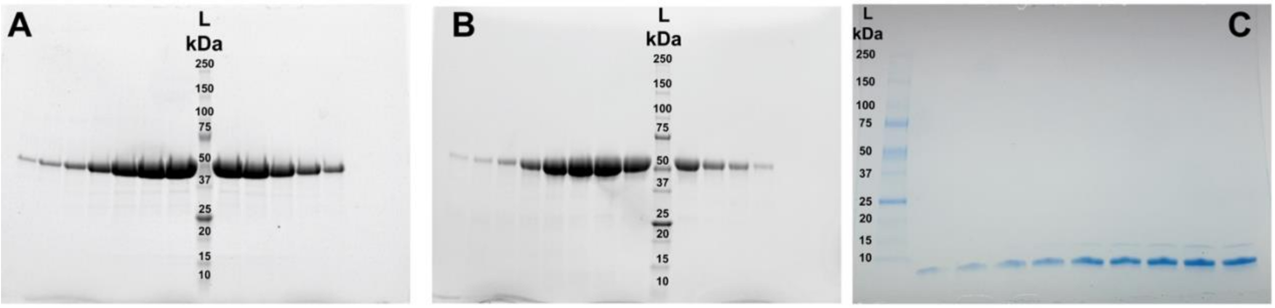

**Figure S1. SDS-Gel of Purified Proteins.**
A, OaaC runs near the 50 kDa marker consistent with its 54 kDa MW.B, OaaA runs slightly above 50 kDa marker consistent with its 58 kDa MW. C, OaaACP runs below the 10 kDa marker consistent with its 9.4 kDa MW.

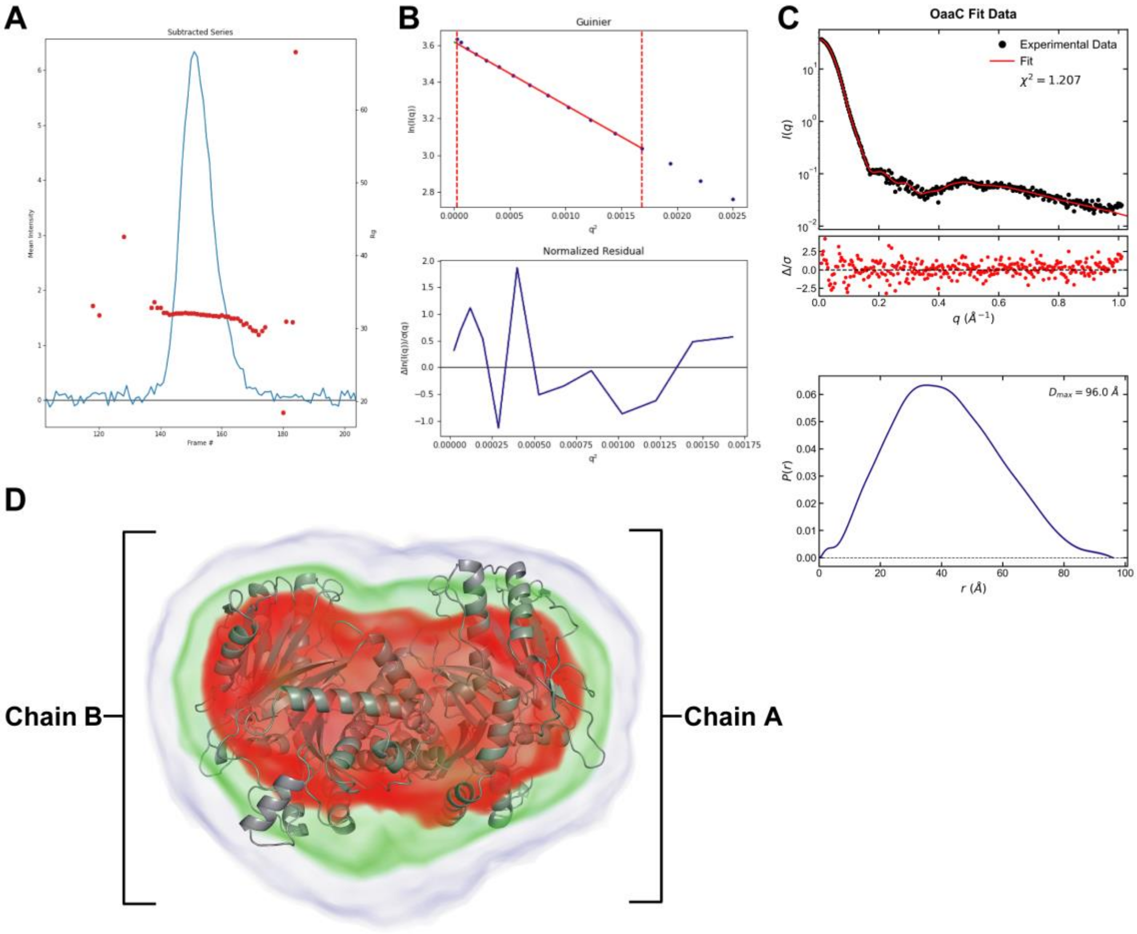

**Figure S2. Analysis of OaaC SEC-SWAXS Data. A,** Scattergram of OaaC, where the mean intensity across each frame of the SEC elution is shown as a blue line (left y-axis). Rg values across the scattergram peak are shown as red circles (right y-axis). B, Guinier analysis of OaaC scattering curve supporting a monodisperse particle in solution (Guinier fit range = 0.16 < q*Rg < 1.31). **C,** DENSS(IFT) fit to the data and resulting P(r) curve indicating a globular particle in solution. **D,** Ab initio DENSS reconstruction using processed OaaC SWAXS profile is shown as a volumetric density map using PyMOL. The map is visualized as a transparent cloud colored according to density ($\sigma$). Lowest contours are rendered in blue (1$\sigma$), intermediate densities in green (5$\sigma$), and the highest density core in red (10$\sigma$). GROMACS-SWAXS model shown in gray cartoon superimposed onto DENSS density map.

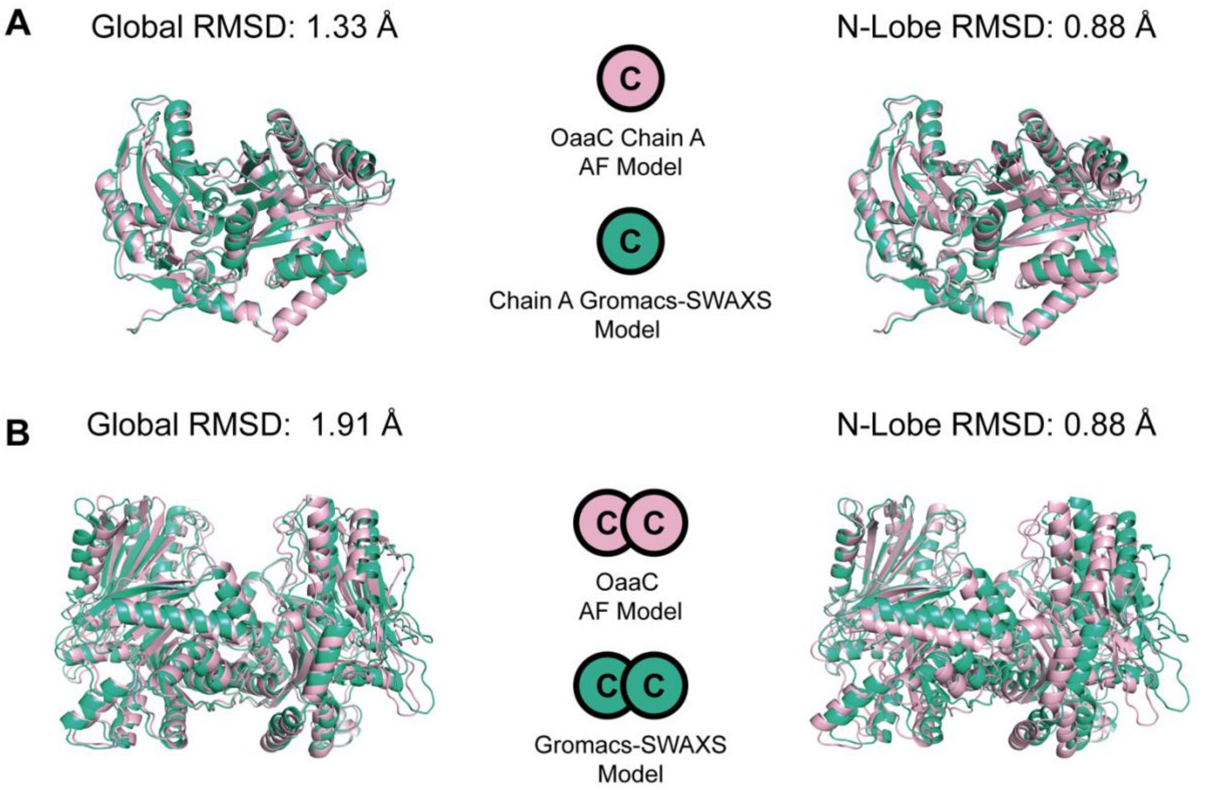

**Figure S3. Structural basis of OaaC solution-state relaxation**. **A,** Superposition of isolated Chain A from the AlphaFold (AF) model (pink) and the representative GROMACS-SWAXS model (teal). Global alignment (left, RMSD = 1.33 Å) and N-lobe anchored alignment (right, RMSD = 0.88 Å) demonstrate that the N-lobe core remains highly stable alongside subtle inter-domain flexibility. **B,** Superposition of the full OaaC dimer. Global alignment (left, RMSD = 1.91 Å) and Chain A N-lobe anchored alignment (right) reveal a quaternary rigid-body shift, demonstrating that the protomers reposition and the solution dimer expands relative to the more compact AF model.

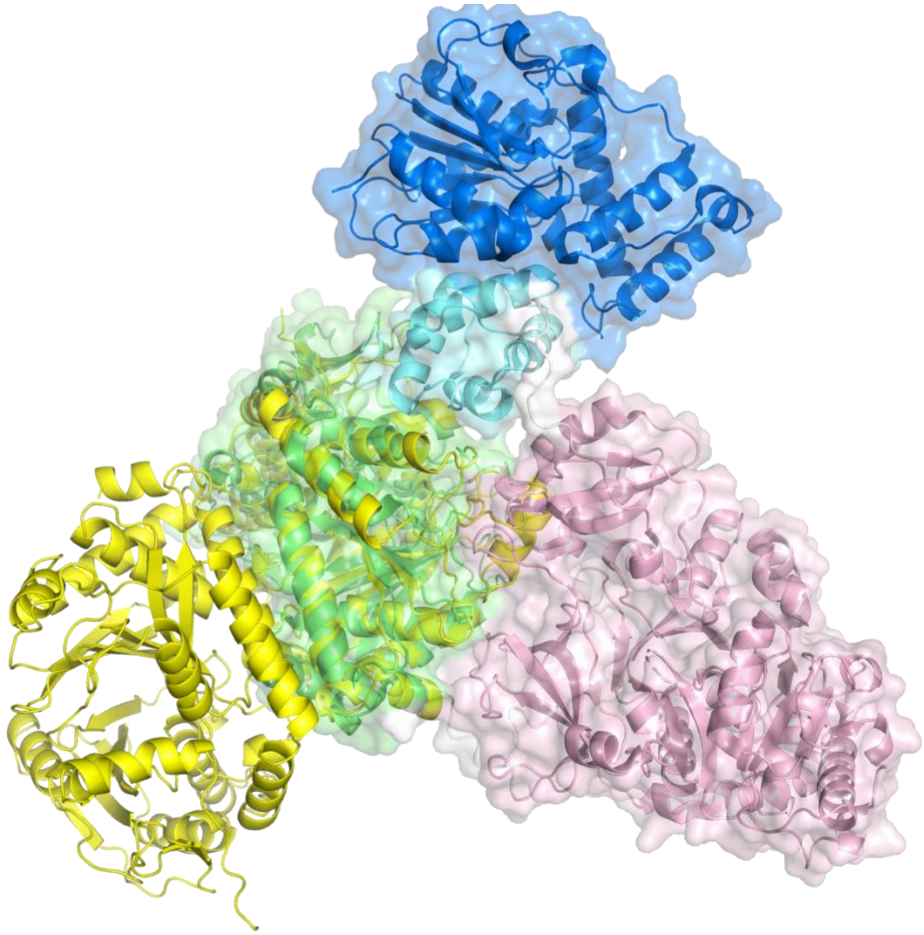

**Figure S4. Comparison of OaaC dimer with an NRPS module.** One chain of the OaaC dimer (yellow) was superimposed on the condensation domain of the AB3403 NRPS module (PDB 4ZXI). The models superimpose with an RMS Displacement of 3.3 Å over 364 Cα atoms. The extensive interface of the AB3403 module between its condensation (green) and adenylation (pink) domains highlights that a different interface is used in the OaaC dimer. The PCP (cyan) and thioesterase (blue) domains of the AB3403 module are also highlighted.

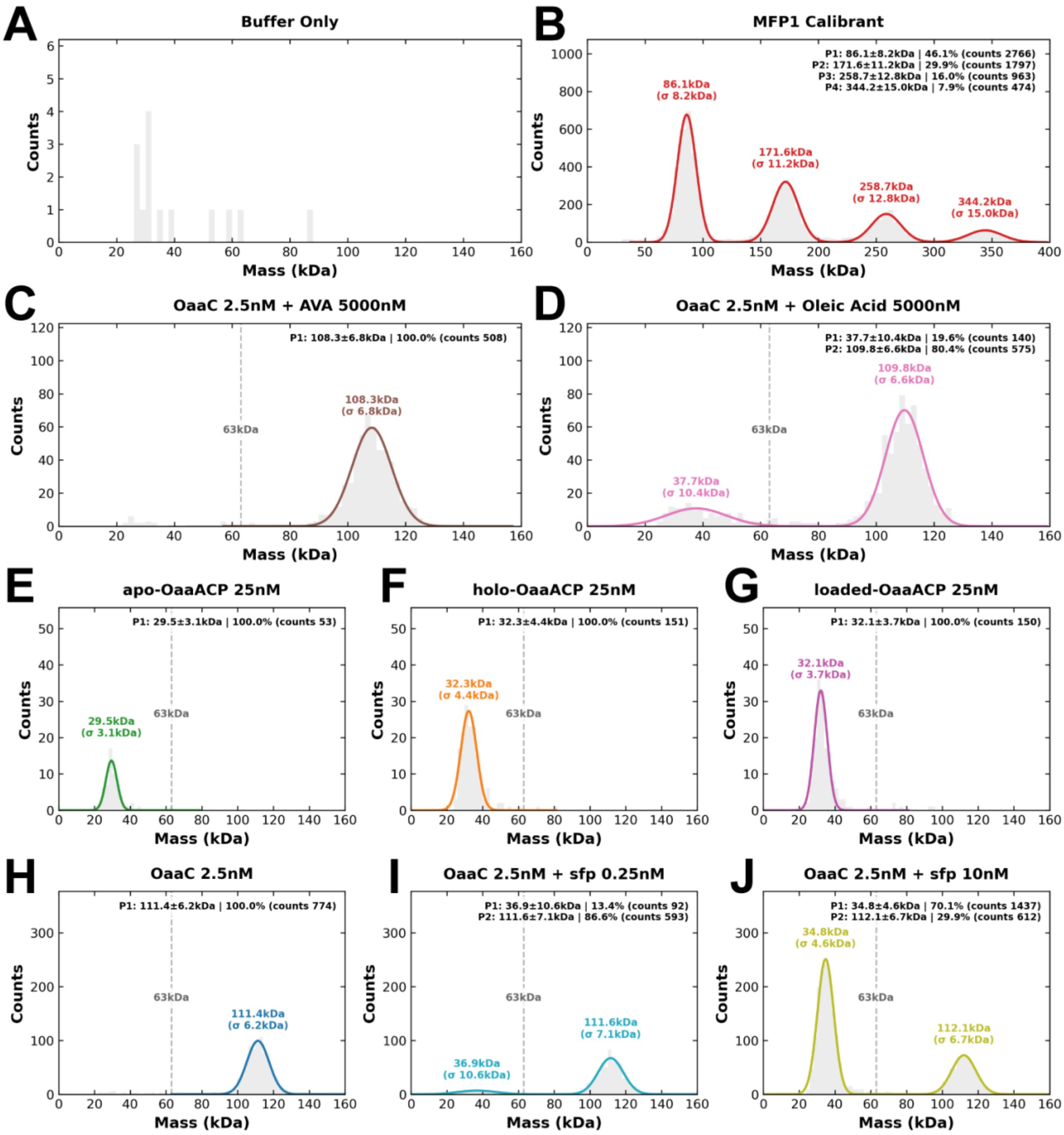

**Figure S5. Mass Photometry of sfp, apo, holo, loaded, OaaACP and OaaC. A,** Clean buffer only slide. **B,** MFP1 calibrant, theoretical MWs: 86kDa, 172kDa, 258kDa, 344kDa. **C,** OaaC at 2.5nM with acceptor substrate aminovaleric acid at 5000nM. **D,** OaaC at 2.5nM with donor substrate oleic acid at 5000nM. **E,** *apo*-OaaACP only at 25nM, MW: 9.4kDa. **F,** *holo*-OaaACP only at 25nM, MW: 9.7kDa. **G,** *loaded*-OaaACP only at 25nM, MW: 10kDa. **H,** OaaC only at 2.5nM. **I,** OaaC at 2.5nM with *sfp* at 0.25nM. **J,** OaaC at 2.5nM with *sfp* at 25nM.

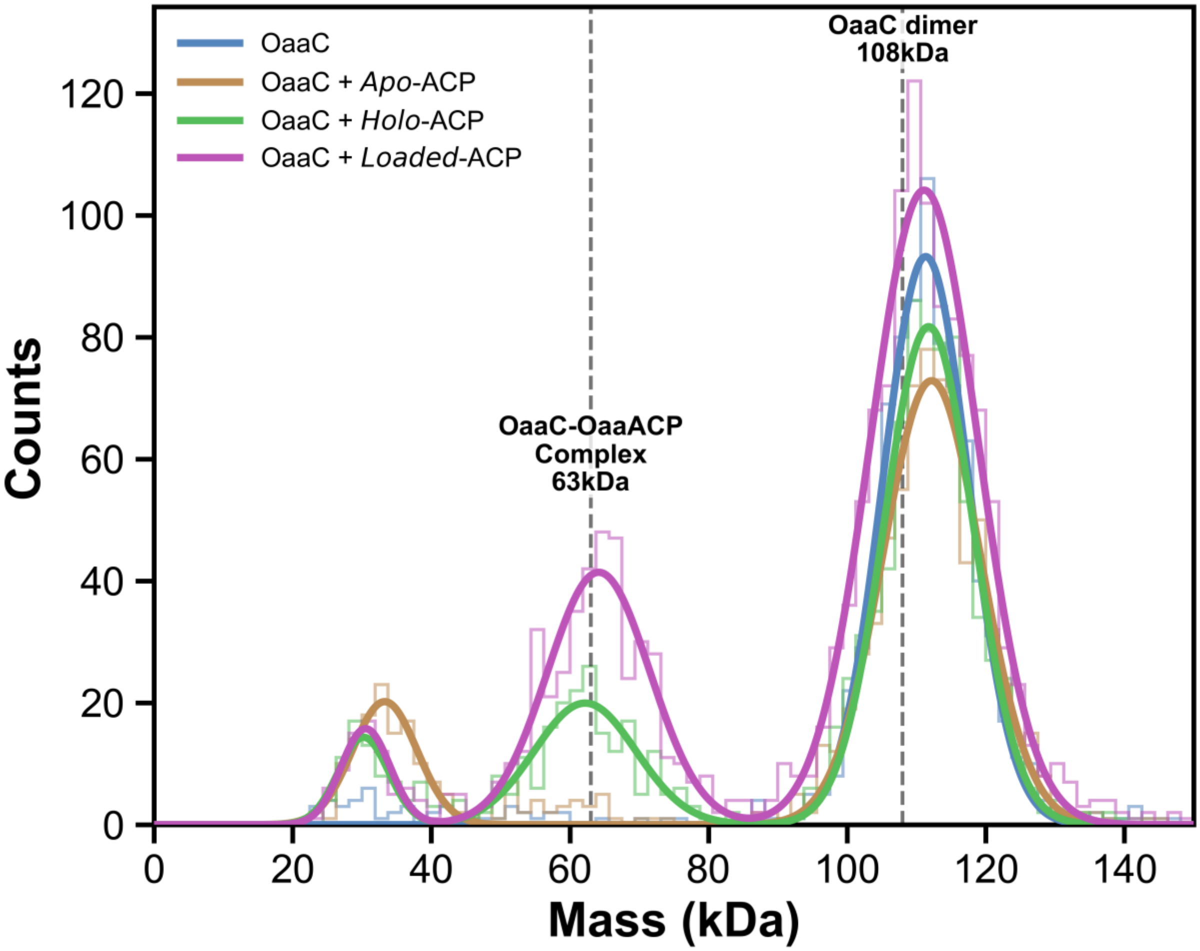

**Figure S6. Mass Photometry Histogram Distribution of sfp, apo, holo, loaded, OaaACP and OaaC.** Overlayed mass photometry raw histogram distributions of OaaC alone and in the presence of a ten-fold excess of OaaACP in distinct functional states (*apo*, *holo*, and acyl-*loaded*). OaaC alone exhibits a dominant species corresponding to the homodimer (111kDa ± 6kDa, counts = 847). Addition of *apo*-OaaACP does not measurably alter the oligomeric distribution, with the homodimer remaining the predominant population (112kDa ± 7kDa, counts = 737). In contrast, incubation with *holo*- and acyl-loaded OaaACP results in retention of the homodimer peak but introduces an additional population centered near ~63 kDa, consistent with formation of an OaaC–OaaACP complex (112kDa ± 6kDa, counts = 725, 62kDa ± 7kDa, counts = 203 and 111kDa ± 8kDa, counts = 1110, 64kDa ± 7kDa, counts = 420 respectively) See SI table 6 for more details. Reference dashed lines denote theoretical molecular weights for the, OaaC–OaaACP complex (63 kDa), and OaaC homodimer (108 kDa).

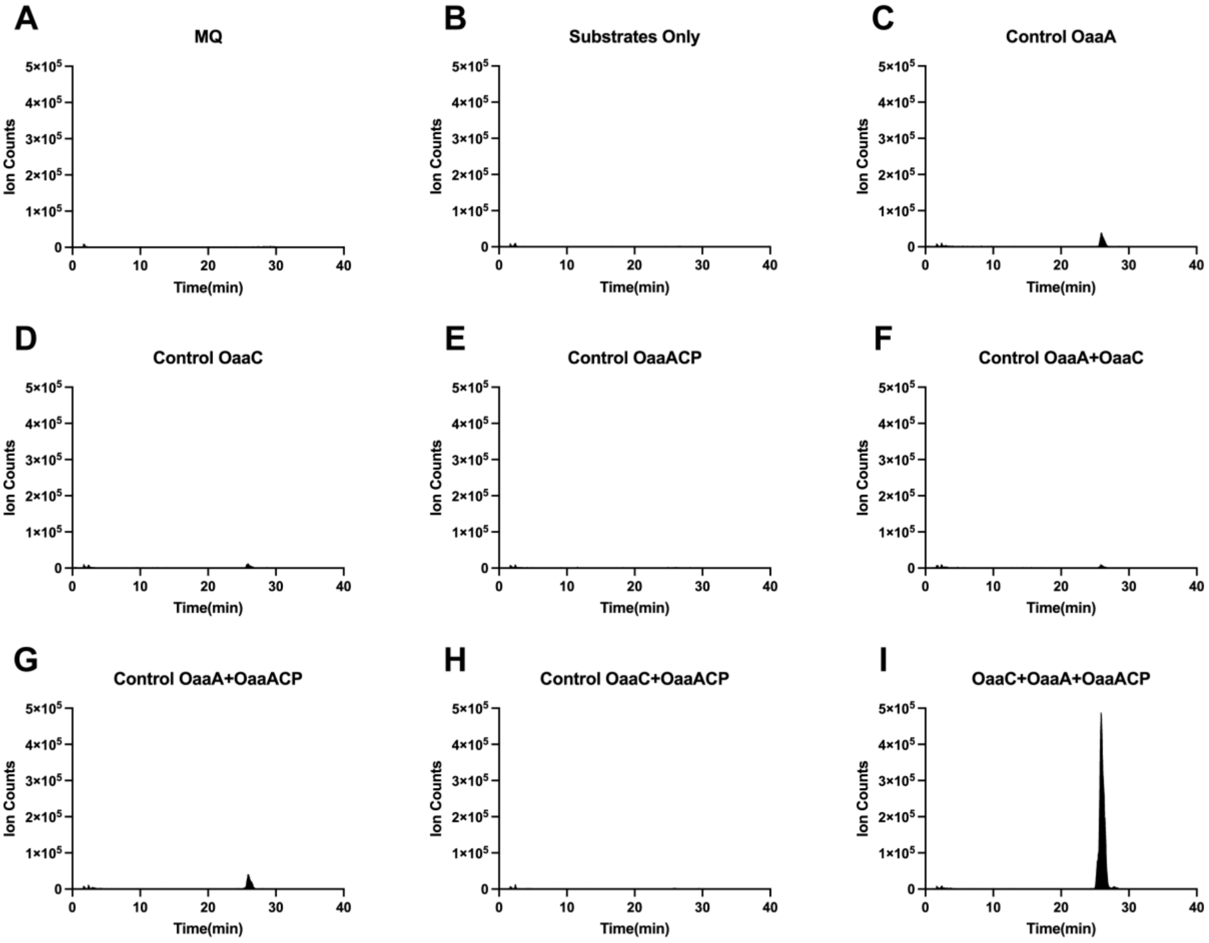

**Figure S7. LC-MS production of oleoyl aminovaleric acid.** Extracted ion chromatograms (EICs) at 382-382.5 m/z depicting oleoyl aminovaleric acid production only in the presence of all three enzymes. **A-H,** each plot represents a different experimental condition where one or more key components for FAA synthesis is removed. **I**, this panel demonstrates that only in the presence of all key enzymes that oleoyl aminovaleric acid synthesis is possible.

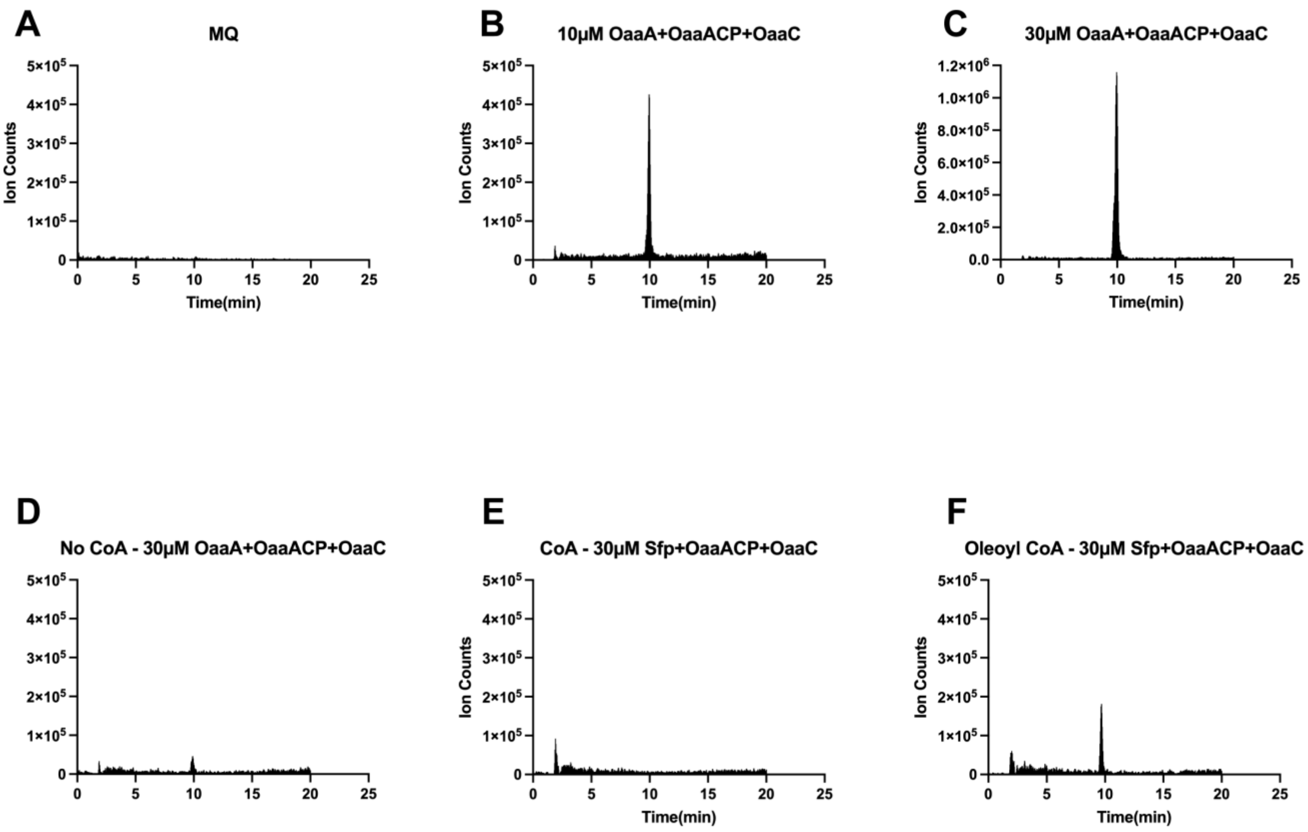

**Figure S8. Oleoyl aminovaleric acid production with Sfp loaded OaaACP. A-C,** EICs at 382-382.5 showing production of FAA with all enzymes included, or **D,** in the absence of CoA. **E,** removing OaaA and using Sfp and CoA leads to no production of FAA. **F,** utilizing *Sfp* and oleoyl-CoA in the absence of OaaA, leads to a one-time loading of OaaACP then enzymatic synthesis of FAA using OaaC and aminovaleric acid.

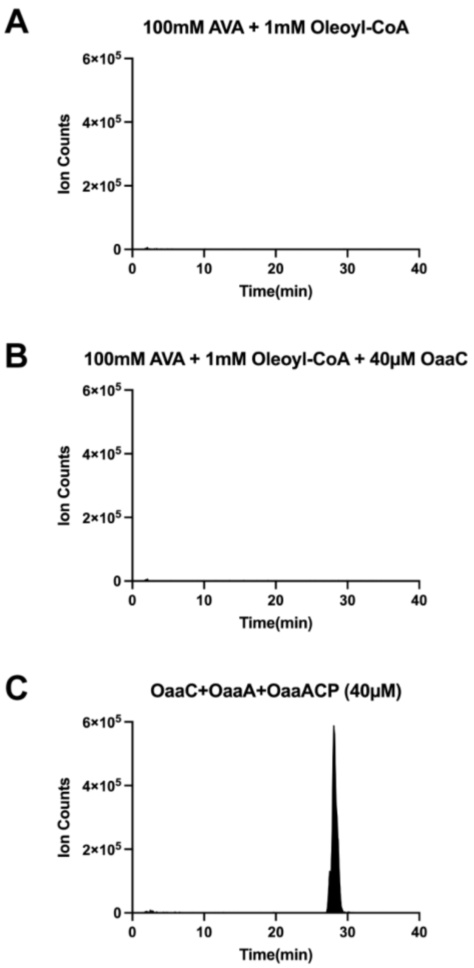

**Figure S9. Assay of OaaC with fatty acyl CoA.** OaaC is unable to use Oleoyl CoA as a donor substrate to form oleoyl aminovaleric. **A,** Substrate only control, in the absence of enzymes no product is formed. **B,** Using OaaC in the presence of AVA and alternative donor substrate oleyl CoA produces no product. **C**, Presence of OaaC, OaaA, OaaACP with AVA and oleic acid produce oleoyl aminovaleric acid. (EICs at 382-382.5)

### Supplemental Tables

**Table S1.** Codon optimized genes used**.**

| Genes | Sequence |
|---|---|
| *ceuT* | CATATGAATAATAATATTACGTTCTTAAATATAGTCGCGGAGTACTGCAACACCCCAGCCGACGAGATAACG<br>AATGACATGCGATTCATAGAGGACTTGGGATTCTCTTCTCTTGATTTCATGACGTTCTTAGGCGACCTTGAG<br>GACACTTTCGACGTAGAGATTAATGAGGACGAAATAATTAATATTCATACAATCGAGGACGCAATAAAGTAC<br>CTTGACAACCTCACGTCCTCAAGTGCCAGTGTGTGATGATGAGCGTCCATAACAGGCGGGCAACAGAAGGGA<br>CGTGGATCCGAATCGAGCTCCGTCGACAAGCTTTAA |
| *ceuC* | CATATGCCGCGTAAATACTATCCGCTGACCCCGAGCCAGAAAATTCATTTTAAACCGATCATTGAATTCGGTACCCAGCAG<br>GTTGCAAATATTAGCATTTGTATGACTCTTCAGGCCCCATTGGATTTTGGACTGCTGAAGAAATGTATCCAGCTGGAATAT<br>GAAAGATATGAATGCCTGCGGATCCGTTTTACTAAAGTGGATCAGAATGGAGAGGTTCGCCAGTATGTCGTAAGTCGTGAT<br>GATCGTGATATTGATTATGAAAATTTAAGTTGGCTTAGCGGCGATGATGCCTATCATCGCATGGAAGAATGGTCACGGATA<br>CCGTTTGATGGTGATAATATACCTATGAACGTGATCAAAATGATTTCCCTGCCAGGCGGTTATAATGGTCTTTATATCAAA<br>ATTGATCACCGGTTAATGGATAGTTGTGGCGCGATCGTTATGGTGAACGATATTATGGAATTATACTGCCACTACAAATTT<br>GGCACTCCATATCCGGAAGATATGGCCTCATTTACGGATATGGTGGAACGTGATCTTAAGAAGTCTACTGATGAGAAAAGA<br>GTGTCGAAAGATCGTATGTATTGGCAGAATGTTTTAGAAGAAAATGGCGAACCCATTTATTCCGATATTCAGGGACAGCGA<br>ATCCTTCAAGAATCACGTCGTCTGCATAATGATAAATCCCTGCGCGCCGCCGATCAAGAAATAAATGATCTGTCGGTCGCG<br>ACCAAGAACTATCATCTTGATGCGGAACCGACGCAAAATCTCCTGGATTTCTGCATGAATAACCATATATCAATGACCAAC<br>CTGATACTCATGGGTATTCGTACATATTTAAGTAAAGCAAATGGCGGTCAGACCGATATCTCAATTCGGAATTATGTCAGT<br>CGTCGGTCTACGCATGCGGAATGGGTTAGCGGTGGTTCACGGGCCATGGCGTATCCGTGTCGAACTATAATCGATCCCGAT<br>ACGGAATTTCTGGATGCGGTTTTCATGATTCAGGATGTGCAGAATCATGTGTATCGGCATTGCAACTATGATCCAGAACTG<br>CTTTCTGATCAGATGAAAGAAATGTTTCATACGCCGCCGCATACTACCTATGAATCAGTTGGGCTGACCTATCAGCCGTTA<br>CCAATTCGGCTGAAGAATCCGCATCTGGAAAACATTTCGGTAAGAAGCATGTGGATCCCGAATGGCACAAGCAAACAGAAA<br>ATATATTTAACCGTTATGCATAGCGCAAATGATCTGGGCCTGAATTTCTATTTTCGGTATCAGACAGCGAGTCTCTCCGAA<br>CAGGACATTGAACTCTTTTACTACTATCTGATGAAAATAATCTTTAAAGGCATTGCTGAACCCGAAATGACCGTGGGTGAG<br>ATTATTGAATGCATATAA |
| *ceuA* | CATATGGAAGAGAACATACTGGAAATCGTCGAGAAATCGTGTCGTATTCATCGCGATGTCATCGCCGTTAAATATCTGTCT<br>CATAGAGAGATCGTGGAGAAAAGTTATGGGGATATGTGGGATGATATACGAAAGACCGCCGTAATTTTACGAAATAATGGT<br>TTATGTGGCACCCATATCGCACTGGTTGGCAGCTCAAGTTATGAATGGATTTGTGCATATATGGCCATTCTGTTTACCGGC<br>AATACAGCAGTGCCCTTGGATGCAAATCTGTCCGTGAGCGAACTTCATGAACTGTTGAATCGGTCAGGGTCCATTGCCCTG<br>TTTTGTGGTGCATCTCGCAAAGATGTGATTACCGAACTTACAGATGATTGTCCGGAAATGAATATCGTTTTCACCATGGAA<br>AAGAAAGTCGACATTGAACATCTGGAAGGTGCAGATAGTAATCCGCAGCTGGCCATTCTGAGCTTTGAACAGCTTCGTAAT<br>GAAATTACCATTCCGGATGATTTTGCGTTCGCGGATCAGGATAAAGATAAAATGTGTACACTGATGTATACATCAGGAACC<br>ACTGGTAAATCAAAAGGTGTGATGCTGTCCCAGTTTAATCTGGCACAGAATGTTGAAAACGTATACGTGAATCTCGAACCT<br>GGAGTAACCATTCTCAGTGTGCTGCCTATTCATCATGCCTTTTGTTTAACCATGGAATGGATGAAAGGAATATCTCTGGGA<br>GCGACAATTTGCATCAATGATTCGTTACTCCACATGCTGAAGAACATGAAACGTTTTCAGCCTGTTGGCATGTTAATGGTT<br>CCACTGATGGTGGAAACCATATACAAGAAACTTAAAGATGTTAATCCGCTGTTGCCCAAGAAATTAGTCGCAAAGGAAGCC<br>TTTGGCGGAAAACTGGAATATATATTTTGCGGCGGTGCCTATCTCGATCCGATGTATGTTACCGAGTTTAAGAAGTATGGC<br>ATCGATATCTTACAAGGGTATGGTATGACAGAATGTTCACCAGTGATATGCAGCAATAATCACCGATATAATAGACCCGGT<br>AGTGTAGGAAAACTGCTTGATAATTGTGCAGTACGTTTTGTCGATGAAGAGATTCAGGTTAAAGGGACGTCCGTTATGAGC<br>GGTTATTATGATATGCCTAACGAAACTGCTGAAGCCTTTCAGGATGGTTGGCTGTGTACAGGAGATTTAGGTTATCTGGAT<br>TCAGATGGGTTTATGTATATAACCGGGCGGAAGAAGAACTTGATTATTCTCGCAAATGGCGAAAACATTTCTCCTGAAGAA<br>CTTGAAGGAAAACTCTCAATCGAACCACTGATTTCAGAAATAGTGATAACCGGGGATGGTAATCATCTCACCGCGCATATT<br>TATCCTGATCAGGATTTCGTAGACAAGAAACACATGGATGCCGCTCGTACGAGCGAAAAGCTGCAGAAAATAATCGACACC<br>TTCAACAAGAATCAGCCAACGTATAAACGTATTTCAGCGTTGGATATTCGTAAAGAGCCATTTGAAAAGTCATCTACAAAG<br>AAGATCAAACGCAACCTGGTCTAA |

**Table S2.** SAS sample details, data collection, analysis, and 3D modelling details for biomolecules in solution.

| | |
|---|---|
| **(*a*) Sample details** | |
| Organism | Bacteria, *Coprococcus eutactus* |
| Source (Catalogue No. or reference) | Condensation domain recombinantly expressed in the BL21-BAP1 cell line. |
| *Scattering particle composition* | |
| Protein(s) | OaaC |
| *Sample environment/configuration* | |
| Solvent composition[d] | 10mM HEPES pH 7.5, 50mM NaCl, 0.2mM TCEP |
| Sample temperature (°C) | ~11°C |
| In beam sample cell[e] | ~11°C |
| Sample injection concentration, mg/ml or g/cm$^3$ | 18 mg/mL |
| Sample injection volume, mL | 28µL |
| SEC column type | 3mL Superdex 200 increase 5/150 |
| SEC flowrate, mL/min | 0.35mL/min |
| **(*b*) SAS data collection** | |
| Data acquisition/reduction software | *py4xs* |
| Source/instrument description or reference | SAXS: Pilatus 1M (1043×981 px); WAXS: Pilatus (981×1043 px) |
| Measured $q$-range ($q_{min} – q_{max}$; Å$^{-1}$, nm$^{-1}$) | 0.005–2.0 Å$^{-1}$ |
| Method for scaling intensities[f] | Arbitrary scale |
| Exposure time(s), number of exposures. For SEC-SAS, final number of sample frames used for averaging. | 2s per frame, 450 frames. Frames 144-161 were averaged and used for downstream analysis. |
| **(*c*) SAS-derived structural parameters** | |
| Methods/Software | RAW/DIFT (DENSS IFT in RAW) |
| *Guinier Analysis* | |
| $I(0) \pm \sigma$ ( a.u) | 37.21±0.03 |
| $R_g \pm \sigma$ (Å) | 32.09±0.04 |
| $min < qR_g < max$ limit (or data point range) | 0.16 – 1.31(1-12) |
| Linear fit assessment (definition)[h] | $R^2$ = 0.9975 |
| *PDDF/P(r) analysis* | |
| $I(0) \pm \sigma$ (a.u.) | 36.95±0.02 |
| $R_g \pm \sigma$ (Å) | 31.66±0.09 |
| $d_{max}$ (Å) | 96 |
| $q$-range (Å$^{-1}$) | 0.005-1.01 |
| $P(r)$ fit assessment (definition)[i] | $\chi^2$ = 1.2 |
| **(*d*) Scattering particle size** | |
| Methods/Software | DENSS (denss-fit-data) |
| *Volume estimates* | |
| Porod volume, $V_p$ (Å$^3$) | 1.83795e+05 ± 3.31653e+03 |
| *Molecular weight (M) estimates (kDa)* | |

| | |
|---|---|
| From chemical composition | 108 kDa |
| From SAS, concentration independent method[j] | 110±1.9kDa |
| **(*e*) Modelling** | |
| *Shape modelling method(s) (if used)* | |
| Software | DENSS |
| $q$-range for fit ($q_{min} - q_{max}$; Å$^{-1}$, nm$^{-1}$) | 0.005-1.01 |
| Symmetry/anisotropy assumptions | N/A |
| Number of individual model reconstructions | 50 |
| $\chi^2$ | 1.653e+00 ± 6.970e-01 |
| Ab initio map resolution (Å) | 47.7 ± 5.6 Å |
| *Atomistic modelling methods* | |
| Software | GROMACS SWAXS |
| $q$-range for fit ($q_{min} - q_{max}$; Å$^{-1}$, nm$^{-1}$) | 0.005-1.01 |
| Number of individual model reconstructions | 200 |
| $\chi^2$ | 1.19 |
| **(*f*) Data and model deposition** | Submitted |
| SASBDB ID | Awaiting review and ID SASBDB assignment |

**Table S3.** Data collection and refinement statistics

| DATA COLLECTION | OAAC |
|---|---|
| PDB CODE | **12KH** |
| Beamline | SSRL |
| Wavelength (Å) | 0.979460 |
| Resolution range (Å) | 51.75 - 2.148 (2.18 - 2.15) |
| Space group | R 3 :H |
| a, b, c (Å) | 122.55 122.55 233.753 |
| **Α, Β, Γ** (°) | 90 90 120 |
| Total reflections | 296740 (10103) |
| Unique reflections | 47625 (1830) |
| Multiplicity | 6.2 (5.5) |
| Completeness (%) | 99.57 (98.59) |
| Mean I/sigma(I) | 8.65 (1.49) |
| $R_{merge}$ | 0.09688 (0.8448) |
| $R_{pim}$ | 0.04176 (0.3916) |
| $CC_{1/2}$ | 0.996 (0.777) |
| **Refinement** | |
| Resolution range (Å) | 51.75 - 2.148 (2.18 - 2.15) |
| Reflections, refinement | 71071 (2664) |
| Reflections, $R_{free}$ | 3601 (145) |
| $R_{work}$ | 0.2015 (0.3689) |
| $R_{free}$ | 0.2386 (0.4037) |
| Protein residues | 925 |
| Ligands of interest | |
| Other molecules | 18 |
| Water molecules | 325 |
| Rms (bonds) (Å) | 0.003 |
| Rms (angles) (°) | 0.59 |
| Rama. Favored (%) | 98.26 |
| Rama. Allowed (%) | 1.41 |
| Rama. Outliers (%) | 0.33 |
| Rotamer outliers (%) | 0.24 |
| Clash score | 3.25 |
| Average B-factor ($Å^2$) | 57.83 |

**Table S4.** Mass photometry statistics and Gaussian mixture modeling for OaaC-ACP complexes. Summary of the statistical parameters derived from Gaussian mixture fitting of mass photometry distributions for OaaC alone and incubated with *Apo*-, *Holo*-, and acyl-*Loaded*-ACP. **Comp**, assigned sub-population component number within the dataset; **μ (kDa)**, the mean mass of the fitted Gaussian peak; **σ (kDa)**, the standard deviation or width of the peak; **Counts**, the estimated number of molecules assigned to the specific component; **%**, the relative fractional abundance of the component calculated from the area under the curve; **N events**, the total number of valid single-molecule scattering events recorded for the entire dataset.

| Dataset | Comp | μ (kDa) | σ (kDa) | Counts | % | N events |
|---|---|---|---|---|---|---|
| OaaC | 1 | 111.3 | 6.2 | 847 | 100.0 | 847 |
| OaaC + *Apo*-ACP | 1 | 33.3 | 4.7 | 135 | 15.5 | 872 |
| OaaC + *Apo*-ACP | 2 | 112.2 | 7.1 | 737 | 84.5 | 872 |
| OaaC + *Holo*-ACP | 1 | 30.2 | 3.6 | 72 | 7.2 | 1000 |
| OaaC + *Holo*-ACP | 2 | 62.2 | 7.3 | 203 | 20.3 | 1000 |
| OaaC + *Holo*-ACP | 3 | 111.8 | 6.4 | 725 | 72.5 | 1000 |
| OaaC + Loaded-ACP | 1 | 30.5 | 3.5 | 76 | 4.7 | 1607 |
| OaaC + Loaded-ACP | 2 | 64.2 | 7.4 | 420 | 26.2 | 1607 |
| OaaC + Loaded-ACP | 3 | 111.1 | 7.8 | 1110 | 69.1 | 1607 |

**Table S5.** Protein sequences

| Protein | Sequence |
|---|---|
| *OaaC*<br>*NCBI Accession*<br>*WP_004853257* | MGSSHHHHHSSGENLYFQGHMPRKYYPLTPSQKIHFKPIIEFGTQQVANISICMTLQAPLDFGLL<br>KKCIQLEYERYECLRIRFTKVDQNGEVRQYVVSRDDRDIDYENLSWLSGDDAYHRMEEWSRIPFD<br>GDNIPMNVIKMISLPGGYNGLYIKIDHRLMDSCGAIVMVNDIMELYCHYKFGTPYPEDMASFTDM<br>VERDLKKSTDEKRVSKDRMYWQNVLEENGEPIYSDIQGQRILQESRRLHNDKSLRAADQEINDLS<br>VATKNYHLDAEPTQNLLDFCMNNHISMTNLILMGIRTYLSKANGGQTDISIRNYVSRRSTHAEWV<br>SGGSRAMAYPCRTIIDPDTEFLDAVFMIQDVQNHVYRHCNYDPELLSDQMKEMFHTPPHTTYESV<br>GLTYQPLPIRLKNPHLENISVRSMWIPNGTSKQKIYLTVMHSANDLGLNFYFRYQTASLSEQDIE<br>LFYYYLMKIIFKGIAEPEMTVGEIIECI |
| *OaaA*<br>*NCBI Accession*<br>*WP_004853261* | MGSSHHHHHSSGENLYFQGHMEENILEIVEKSCRIHRDVIAVKYLSHREIVEKSYGDMWDDIRKT<br>AVILRNNGLCGTHIALVGSSSYEWICAYMAILFTGNTAVPLDANLSVSELHELLNRSGSIALFCG<br>ASRKDVITELTDDCPEMNIVFTMEKKVDIEHLEGADSNPQLAILSFEQLRNEITIPDDFAFADQD<br>KDKMCTLMYTSGTTGKSKGVMLSQFNLAQNVENVYVNLEPGVTILSVLPIHHAFCLTMEWMKGIS<br>LGATICINDSLLHMLKNMKRFQPVGMLMVPLMVETIYKKLKDVNPLLPKKLVAKEAFGGKLEYIF<br>CGGAYLDPMYVTEFKKYGIDILQGYGMTECSPVICSNNHRYNRPGSVGKLLDNCAVRFVDEEIQV<br>KGTSVMSGYYDMPNETAEAFQDGWLCTGDLGYLDSDGFMYITGRKKNLIILANGENISPEELEGK<br>LSIEPLISEIVITGDGNHLTAHIYPDQDFVDKKHMDAARTSEKLQKIIDTFNKNQPTYKRISALD<br>IRKEPFEKSSTKKIKRNLV |
| *OaaACP*<br>*NCBI Accession*<br>*WP_004853259* | MGSSHHHHHSSGENLYFQGHMNNNITFLNIVAEYCNTPADEITNDMRFIEDLGFSSLDFMTFLGD<br>LEDTFDVEINEDEIINIHTIEDAIKYLDNLTSSSASV |

Sequence underlined represents N-5x-His-tag and TEV cleavage site residues (ENLYFQ^G) contributed by vector pET28b.
